# SMARCAD1 Mediated Active Replication Fork Stability Maintains Genome Integrity

**DOI:** 10.1101/2020.10.05.326223

**Authors:** Calvin Shun Yu Lo, Marvin van Toorn, Vincent Gaggioli, Mariana Paes Dias, Yifan Zhu, Eleni Maria Manolika, Wei Zhao, Marit van der Does, Chirantani Mukherjee, João G S C Souto Gonçalves, Martin E van Royen, Pim J French, Jeroen Demmers, Ihor Smal, Hannes Lans, David Wheeler, Jos Jonkers, Arnab Ray Chaudhuri, Jurgen A Marteijn, Nitika Taneja

**Author notes:** These authors contributed equally to this work (Marvin van Toorn, Vincent Gaggioli).

## Abstract

Stalled fork protection pathway mediated by BRCA1/2 proteins is critical for replication fork stability that has implications in tumorigenesis. However, it is unclear if additional mechanisms are required to maintain replication fork stability. We describe a novel mechanism by which the chromatin remodeler SMARCAD1 stabilizes active replication forks that is essential for resistance towards replication poisons. We find that loss of SMARCAD1 results in toxic enrichment of 53BP1 at replication forks which mediates untimely dissociation of PCNA via the PCNA-unloader, ATAD5. Faster dissociation of PCNA causes frequent fork stalling, inefficient fork restart and accumulation of single-stranded DNA resulting in genome instability. Although, loss of 53BP1 in SMARCAD1 mutants restore PCNA levels, fork restart efficiency, genome stability and tolerance to replication poisons; this requires BRCA1 mediated fork protection. Interestingly, fork protection challenged BRCA1-deficient naïve- or PARPi-resistant tumors require SMARCAD1 mediated active fork stabilization to maintain unperturbed fork progression and cellular proliferation.

## INTRODUCTION

Most BRCA-mutated cancers acquire resistance towards chemotherapeutic agents such as cisplatin and PARP inhibitors (PARPi) (*1*). At present, besides the restoration of homologous recombination (HR), loss of PAR glycohydrolase (PARG) or acquired protection of stalled replication forks provides a mechanism that can promote drug resistance in BRCA-deficient genetic background (*1-4*). However, identification of additional mechanisms underlying resistance to chemotherapeutics can provide a real opportunity to improve therapies in BRCA-deficient cancer patients.

BRCA-proteins play a genetically separable role at the site of double-stranded breaks (DSBs) where they mediate an error-free HR repair and at replication forks where they facilitate protection of reversed forks from extensive nuclease-mediated degradation, to maintain genome stability (*2, 3, 5-7*). Similarly, the factors of non-homologous end joining (NHEJ), an error-prone pathway, along with their role in repair of DSBs have been shown to associate with stalled forks either for their protection or to promote their restart (*2, 8, 9*). However, the factors involved in limiting fork stalling and subsequent restarting of forks upon endogenous or exogenously induced replication stress are poorly understood.

Proliferating Cell Nuclear Antigen (PCNA) is a DNA clamp that associates with the active replication forks and functions as a processivity factor for DNA polymerases to carry out the DNA synthesis process but dissociates from stalled forks via an active unloading mechanism (*8, 10-12*). During replication, PCNA rings are repeatedly loaded and unloaded by the replicating clamp loader replication factor C (RFC) complex (*13*) and an alternative PCNA ring opener, ATAD5 (ELG1 in yeast)-replication factor C-like complex (ATAD5-RLC). ATAD5-RLC unloads replication-coupled PCNA after ligation of Okazaki fragment and termination of DNA replication (*14-16*). Maintenance of the delicate balance of PCNA levels onto DNA is crucial since PCNA levels can influence chromatin integrity (*17-19*) and persistent PCNA retention on DNA causes genome instability (*20-22*). However, mechanisms by which PCNA levels are regulated on replicating chromatin and the factors involved in this process, still remain elusive.

Here we uncover a novel function of human SMARCAD1 in regulating the fine control of PCNA levels at forks, which is required for the maintenance of replication stress tolerance and genome stability. SMARCAD1, a DEAD/H box helicase domain protein, belongs to a highly conserved ATP-dependent SWI/SNF family of chromatin remodelers. ATPase remodeling activity of SMARCAD1 is crucial for its function in HR repair as well as in maintenance of histone methyl marks for re-establishment of heterochromatin (*23, 24*).

In this study, we generated a separation-of-function mutant of human SMARCAD1, efficient in its HR function but defective in its interaction with the replication machinery. This strategy led to uncover a previously unrecognized role of SMARCAD1 in maintaining stability of active (unperturbed and restarted) replication forks, which is responsible for mediating resistance towards replication poisons. In the absence of SMARCAD1, replication fork progression requires BRCA1 to maintain the integrity of stalled forks to allow their restart. Furthermore, SMARCAD1 maintains replication fork stability and cellular viability in BRCA1 deficient naïve or chemoresistant mouse breast tumor-organoids, highlighting its essential role in the survival of tumor cells. Our results suggest a conserved role of SMARCAD1 and BRCA1 proteins at replication forks, SMARCAD1 at active forks while BRCA1 at stalled forks, to safeguard replication fork integrity and ensure genome stability.

## RESULTS

### SMARCAD1 is preferentially enriched at unperturbed replication forks

Most factors associated with the active replisome are required to maintain the stability of the replication forks and could also be important for mediating efficient restart after stalling. In order to specifically identify novel factors involved in the stability of unperturbed forks, we performed isolation of Proteins On Nascent DNA (iPOND) coupled to Stable Isotope Labeling with Amino acids in Cell culture (SILAC)-based quantitative mass-spectrometry (*8, 25*). Mouse embryonic stem cells (mESC) were used to compare the proteins present at unperturbed active replication forks vs hydroxyurea (HU)-induced stalled replication fork (fig. S1A). In total 1443 common proteins were identified from two independent experiments (fig. S1, B and C). Consistent with previous reports, we observed a greater than two-fold increase in replication stress response proteins, including RAD51 and BRCA1, at stalled forks (Fig. 1A) (*8, 25*). Levels of core components of the replicative helicase, such as MCM6, remained largely unchanged during early replication stress (Fig. 1A). As shown previously (*8*), PCNA was enriched ∼2-fold at the unperturbed forks when compared to the stalled forks, confirming that PCNA associates preferentially with active forks and showing proof-of-principle of this approach (Fig. 1A and fig. S1B). Among 66 proteins showing preferential enrichment at unperturbed replication forks (Fig. S1C), we identified SMARCAD1, a conserved SWI/SNF chromatin remodeler (Fig. 1A and fig. S1B). Interestingly, KAP1/ TRIM28, a previously reported SMARCAD1 interacting partner, showed no preferential enrichment, a behavior that is similar to that of the MCM6 helicase, suggesting an additional and independent role of SMARCAD1 in replication fork dynamics (Fig. 1A) (*8*).

**Fig. 1.**
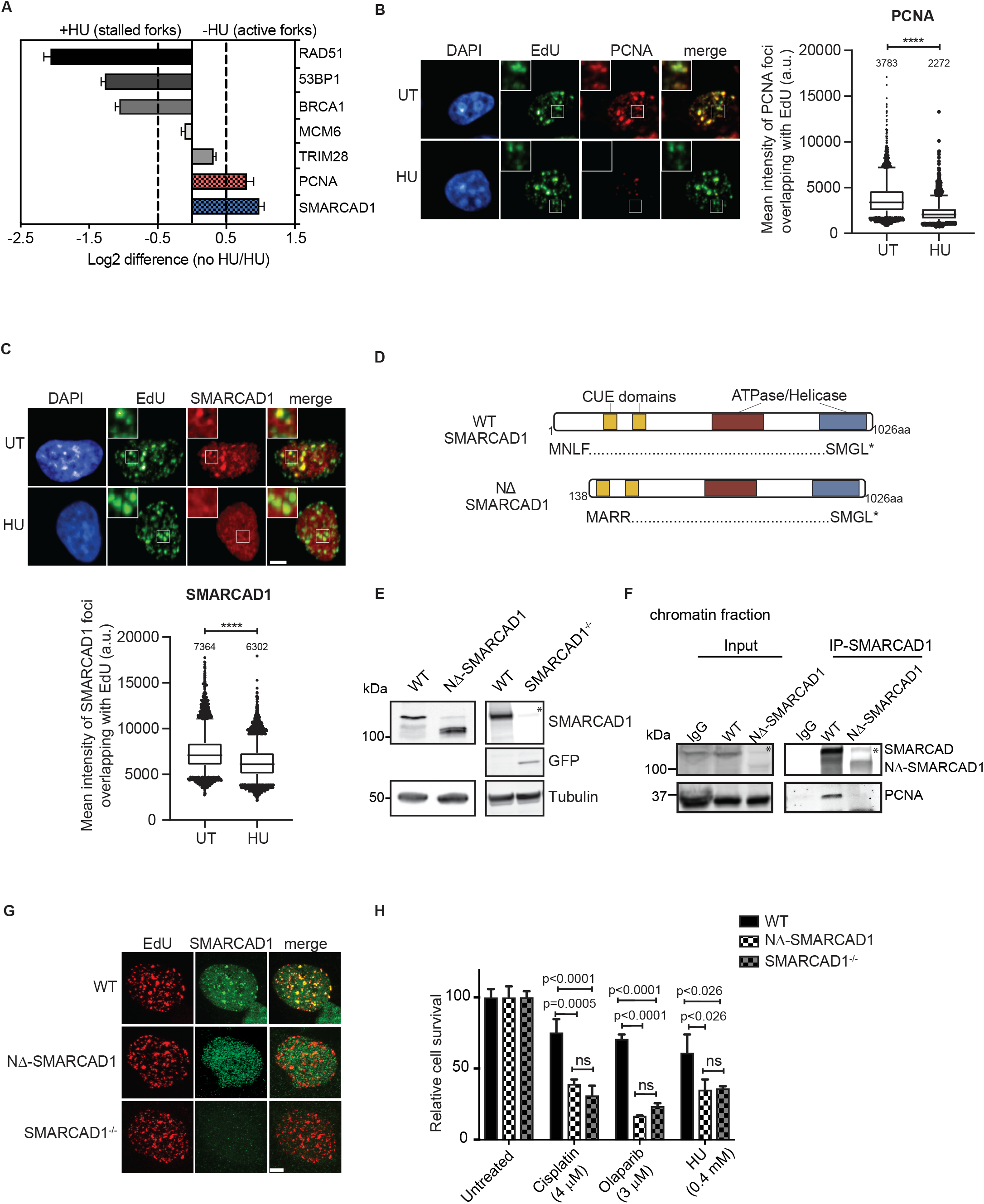
PCNA-interacting domain of SMARCAD1 is required for its localization to active replication forks. (**A**) Bar graph showing fold upregulation of selected proteins in unperturbed (no HU) and HU treated conditions based on their SILAC H:L ratios. (**B**) Left: Representative high-content microscopy images showing the co-localization of chromatin bound PCNA (red) to the sites of DNA replication marked with EdU (green) in the presence or absence of HU in WT cells (note that for the HU condition EdU labelling was performed prior to HU treatment). Right: Quantification of mean intensity of PCNA foci overlapping with EdU are shown as box plot. n>3000 cells with EdU foci per condition were analysed in mid-late S phase cells. Numbers above each scatter plot indicate the mean intensity of each PCNA foci overlapping with EdU. (\*\*\*\**P* ≤ 0.0001, unpaired *t*-test). (**C**) Top panel: Representative high-content microscopy images showing the co-localization of chromatin bound SMARCAD1 (red) to the sites of DNA replication marked with EdU (green) in the presence or absence of HU in WT cells (note that for the HU condition EdU labelling was performed prior to HU treatment) (scale bar = 5μm). Quantification of mean intensity of SMARCAD1 foci overlapping with EdU are shown as box plot. n>3000 cells with EdU foci per condition were analysed in mid-late S phase cells. Numbers above each scatter plot indicate the mean intensity of each SMARCAD1 foci overlapping with EdU. (\*\*\*\**P* ≤ 0.0001, unpaired *t*-test). (**D**) Schematic overview of the protein domains in full-length SMARCAD1 and NΔ-SMARCAD1. (**E**) Immunoblot showing SMARCAD1 levels in WT, NΔ-SMARCAD1 and SMARCAD1^-/-^ MRC5 cells. Tubulin is used as a loading control. (* represents a non-specific band, confirmed by lack of full-length transcripts in SMARCAD1^-/-^, as shown in Fig. S1F) (**F**) Crosslinked immunoprecipitation of SMARCAD1 was performed in WT and NΔ-SMARCAD1 cells using SMARCAD1 antibody. Western blots were performed using antibodies against PCNA and SMARCAD1 (* represents a non-specific band). (**G**) Representative images showing expression of SMARCAD1 (green) and EdU (red) in WT, NΔ-SMARCAD1 and SMARCAD1^-/-^ cells (scale bar = 5μm). Note that NΔ-SMARCAD1 protein associates with chromatin but does not colocalize with EdU signal unlike in WT-SMARCAD1. (**H**) Quantification of colony survival assay in WT, NΔ-SMARCAD1 and SMARCAD1^-/-^ cells treated with HU, cisplatin and olaparib. HU was given for 48 hours before release for 6 extra days. The mean and S.D. from three independent experiments is represented. (ns, non-significant, Unpaired t-test)

To confirm our iPOND-SILAC-MS data and to assess if the preferential enrichment of SMARCAD1 and PCNA at unperturbed replication forks is conserved across species, we performed immunofluorescence assays to measure the localization of these proteins with respect to the sites of replication in MRC5 human fibroblast cells. Sites of active DNA replication were labelled with EdU, and the localization of the chromatin-bound fraction of SMARCAD1, PCNA and RAD51 within the sites of replication was measured in the presence or the absence of hydroxyurea (HU) using single-cell based, high-content microscopy. Consistent with the results of iPOND-SILAC-MS in mESCs, we observed that chromatin bound SMARCAD1 and PCNA foci specifically colocalized with EdU. However, upon HU treatment both these proteins showed a significant decrease in intensity at replication sites, suggesting that both SMARCAD1 and PCNA associate with unperturbed replication forks but dissociate from stalled forks (Fig. 1, B and C). As expected, RAD51 was found to be enriched significantly at replication sites upon HU treatment, suggesting a positive enrichment at stalled forks in contrast to PCNA and SMARCAD1 (fig. S1D) (*8, 25*).

### NΔ-SMARCAD1 lacks PCNA interaction and thereby, association with replication forks

The N-terminal region of SMARCAD1 has been shown to be responsible for the PCNA-mediated localization of SMARCAD1 to replication forks (*24, 26*). To explore the role of this interaction at replication forks, we generated a SMARCAD1 mutant, using MRC5 cells, in which the canonical start site is disrupted, and translation begins downstream at the next available start codon (Fig. 1D). Expression of this mutant gene results in a 137 amino acids N-terminally truncated product, designated as NΔ-SMARCAD1 that lacks the region responsible for its interaction with PCNA (*26*). The NΔ-SMARCAD1 protein is approximately 100 kDa in size (Fig. 1E) and retains the downstream CUE1, CUE2, ATPase and Helicase domains (fig. S1E), crucial for chromatin remodeling and DNA repair functions (*24, 27*), intact. For comparative analysis, we also generated a complete SMARCAD1 knockout (SMARCAD1^-/-^) by replacing the SMARCAD1 gene with a mClover (a GFP variant) reporter gene (Fig. 1E). Both qRT-PCR assays of the SMARCAD1 coding region as well as RNASeq-based transcriptome analysis of cells containing the full length (WT) and those containing the truncated form (NΔ-SMARCAD1) confirmed that expression levels of the two SMARCAD1 alleles were nearly identical (fig. S1, E and F). As expected, cells containing the knockout, SMARCAD1^-/-^, showed a lack of transcripts specific to the coding region of the gene.

To test the interaction between PCNA and the NΔ-SMARCAD1 mutant, we generated a heterogeneously expressed GFP-tagged PCNA allele in both WT and NΔ-SMARCAD1 genetic backgrounds (fig. S1G). Crosslinked chromatin immunoprecipitation of GFP-tagged PCNA confirmed that even though NΔ-SMARCAD1 associates with chromatin, it did not interact with GFP-PCNA, whereas the full-length wildtype SMARCAD1 protein retains this interaction (fig. S1H) as previously reported (*24*). Similarly, reverse chromatin immunoprecipitation of WT-SMARCAD1 and NΔ-SMARCAD1 protein confirmed the lack of interaction between PCNA and NΔ-SMARCAD1 protein (Fig. 1F). To determine whether a SMARCAD1 interaction with PCNA is required for its association with replication sites, we performed an immunofluorescence analysis to measure the localization of SMARCAD1 mutants at sites of DNA replication marked with EdU. Our data show that chromatin bound foci of full length SMARCAD1 colocalized with EdU positive sites as previously reported (*24*) (Fig. 1G). As expected, no specific SMARCAD1 signal could be seen in SMARCAD1 knockout (SMARCAD1^-/-^) cells. Consistent with our crosslinked IP data (Fig. 1F and fig. S1H), NΔ-SMARCAD1 showed nuclear localization but no colocalization with EdU signals (Fig. 1G), suggesting that NΔ-SMARCAD1 associates with chromatin but is not enriched at sites of replication.

### Role of SMARCAD1 at the replication fork and not in HR, mediates tolerance to replicative stress

Next, we sought to determine if loss of SMARCAD1 association with replication forks affects cellular resistance to fork stalling agents such as hydroxyurea (HU), cisplatin or the PARP inhibitor, olaparib. Both NΔ-SMARCAD1 and SMARCAD1^-/-^ cells showed significant sensitivity to the replication poisons, suggesting that the presence of SMARCAD1 at replication forks is crucial for resistance to replication stress (Fig. 1H). To further explore the role of SMARCAD1 during DNA replication, we analyzed S phase progression by measuring EdU incorporation using high-content microscopy. We imaged >2000 cells and plotted for quantitative image-based cytometry analysis (QIBC) to obtain single-cell based cell cycle profile (*28*). Both NΔ-SMARCAD1 and SMARCAD1^-/-^ cells displayed reduction in EdU intensities relative to WT cells suggesting loss of SMARCAD1 at forks causes DNA replication defects (fig. S1I).

Since the loss of SMARCAD1 causes defects in HR repair of DSBs due to inefficient DNA end-resection (*23, 27, 29*), we next tested whether cells expressing NΔ-SMARCAD1 also exhibited defects in HR repair. We measured HR efficiency using a DR-GFP reporter assay (*30*). Remarkably, NΔ-SMARCAD1 cells had an HR efficiency similar to that of WT (Fig. 2A). However, HR efficiency was significantly reduced in both, WT and NΔ-SMARCAD1 cells when SMARCAD1 was knocked down in these cells using siRNA, similar to that observed for BRCA1 knockdown (Fig. 2A). These data suggest that, although the complete loss of SMARCAD1 results in defective HR, expression of the truncated NΔ-SMARCAD1 retains HR proficiency. Additionally, chromatin fractionation and observation of RAD51 focus formation by immunofluorescence using high content microscopy, both showed a remarkable increase in chromatin-bound RAD51 upon olaparib treatment in both WT and NΔ-SMARCAD1, but not in SMARCAD1 deficient cells (Fig. 2, B and C). This data further confirms that NΔ-SMARCAD1 cells are proficient in the loading of RAD51 in response to DNA damage unlike SMARCAD1^-/-^. Surprisingly however, both the mutants show similar sensitivity towards drugs causing replication stress, olaparib, cisplatin and HU (Fig. 1H, 2D and fig. S2A), arguing in favor of an uncoupling between HR repair function and resistance to replication stress in the NΔ-SMARCAD1 cells, corroborating it to be a separation-of-function mutant.

**Fig. 2.**
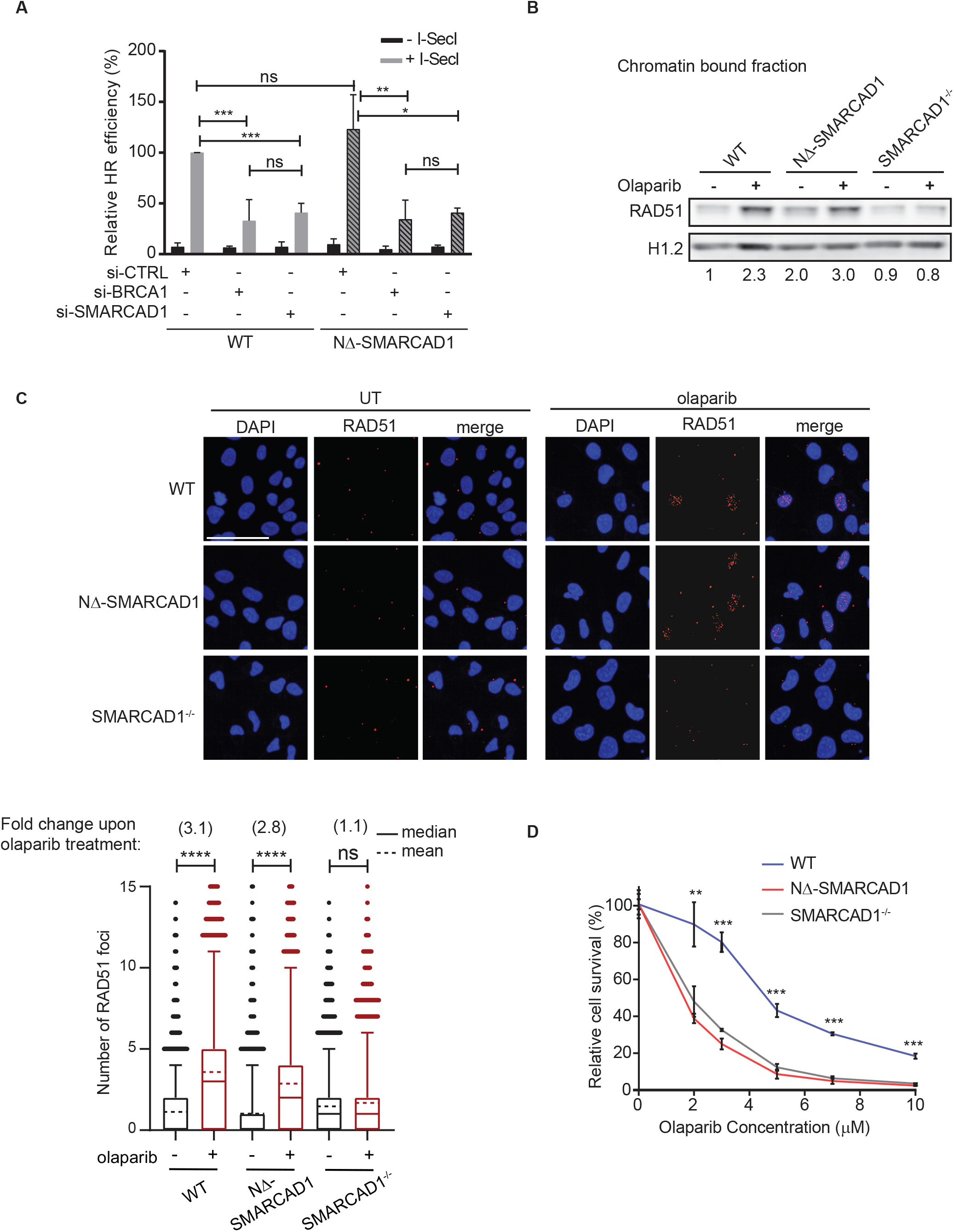
SMARCAD1 provides resistance towards replication poisons, independent of its role in HR repair pathway. (**A**) Quantification of HR efficiency using DR-GFP reporter assay. DR-GFP reporter and pcBASceI constructs were co-transfected into WT and NΔ-SMARCAD1 MRC5 cells. Relative HR efficiency representing the percentage of GFP positive cells normalised to transfection efficiency of the respective cell line is plotted. The mean and S.D. from three independent experiments is represented. (\*\*\**P* ≤ 0.001,\*\**P* ≤ 0.01, \**P* ≤ 0.05,ns, non-significant, Unpaired *t*-test) (**B**) Immunoblot showing the chromatin bound fraction of RAD51 upon 7μM olaparib treatment for 24 hours in WT, NΔ-SMARCAD1 and SMARCAD1^-/-^ cells. H1.2 is used as a loading control. The numbers below the blots show the fold change of RAD51 after normalisation with H1.2 as compared to WT untreated samples, for the given blot (total n = 3). (**C**) Top: Representative high content microscopy images depicting RAD51 foci formation upon 7μM olaparib treatment for 24 hours in WT, NΔ-SMARCAD1 and SMARCAD1^-/-^ cells. Scale bar = 50 μm. Bottom: Quantification of the number of RAD51 foci upon 7μm olaparib treatment for 24 hours using high-content microscopy. 4700 cells were analyzed in each condition. Solid line and dotted line represent median and mean, respectively. (\*\*\*\**P* ≤ 0.001, ns, non-significant, One-way ANOVA). Number above represented the fold change of RAD51 foci upon olaparib treatment compared to its own untreated samples. (**D**) Quantification of colony survival assay in WT, NΔ-SMARCAD1 and SMARCAD1^-/-^ cells treated with different concentrations of olaparib. Error bars stand for ±S.D. (n=3). (\*\*\**P* ≤ 0.001,\*\**P* ≤ 0.01, unpaired *t*-test)

We also performed transcriptome analysis to test whether the drug sensitivity observed in SMARCAD1 mutant cells could be a result of transcription deregulation of DDR genes in these cells, since transcription may be affected by its chromatin-remodeling role. We observed a mild dysregulation in a subset of non-DDR genes (≥ 1.5 fold change in expression) in either NΔ-SMARCAD1 or SMARCAD1^-/-^ cells whereas almost no anomalous expression was observed in either mutant for a set of DDR genes (N=179) (*31*), that included both HR and NHEJ DNA damage response genes (fig. S2B). This suggests that the function of SMARCAD1 in promoting drug tolerance is unrelated to its role in heterochromatin maintenance or in transcriptional regulation. Furthermore, the efficient loading of RAD51 and the HR proficiency of cells expressing NΔ-SMARCAD1, in contrast to those lacking SMARCAD1, is most likely not due to a differential transcriptome or cell cycle profile but due to the presence of intact CUE and ATPase-Helicase domains in NΔ-SMARCAD1 that are essential for its HR function (*23, 29*). Intriguingly, the loss of PCNA interaction and association with the fork is the main cause for SMARCAD1 depleted cells to show sensitivity towards replication stress inducing drugs.

### SMARCAD1 facilitates normal replication fork progression and efficient restart upon replication stress

SMARCAD1 mutants displayed moderate but significant defects in progression through S phase (fig. S1I). To further monitor the dynamics of individual replication forks we performed DNA fiber assay. We sequentially labeled WT and SMARCAD1 mutants (NΔ-SMARCAD1 and SMARCAD1^-/-^) cells with CldU (red) and IdU (green), followed by track length analysis. Interestingly, NΔ-SMARCAD1 cells exhibited a significant difference in the track lengths of both CldU and IdU in comparison to WT but similar to SMARCAD1^-/-^ cells (Fig. 3A). To test the possibility that accumulation of DNA damage over time in the mutant cells was causing the replication fork defect observed, we also analyzed fork progression in cells in which SMARCAD1 was depleted transiently with siRNA. The transient knock down of SMARCAD1 resulted in similar fork progression defects than the one observed in NΔ-SMARCAD1 and SMARCAD1^-/-^ (Fig. 3A). This suggests that SMARCAD1 directly facilitates the progression of replication forks.

**Fig. 3.**
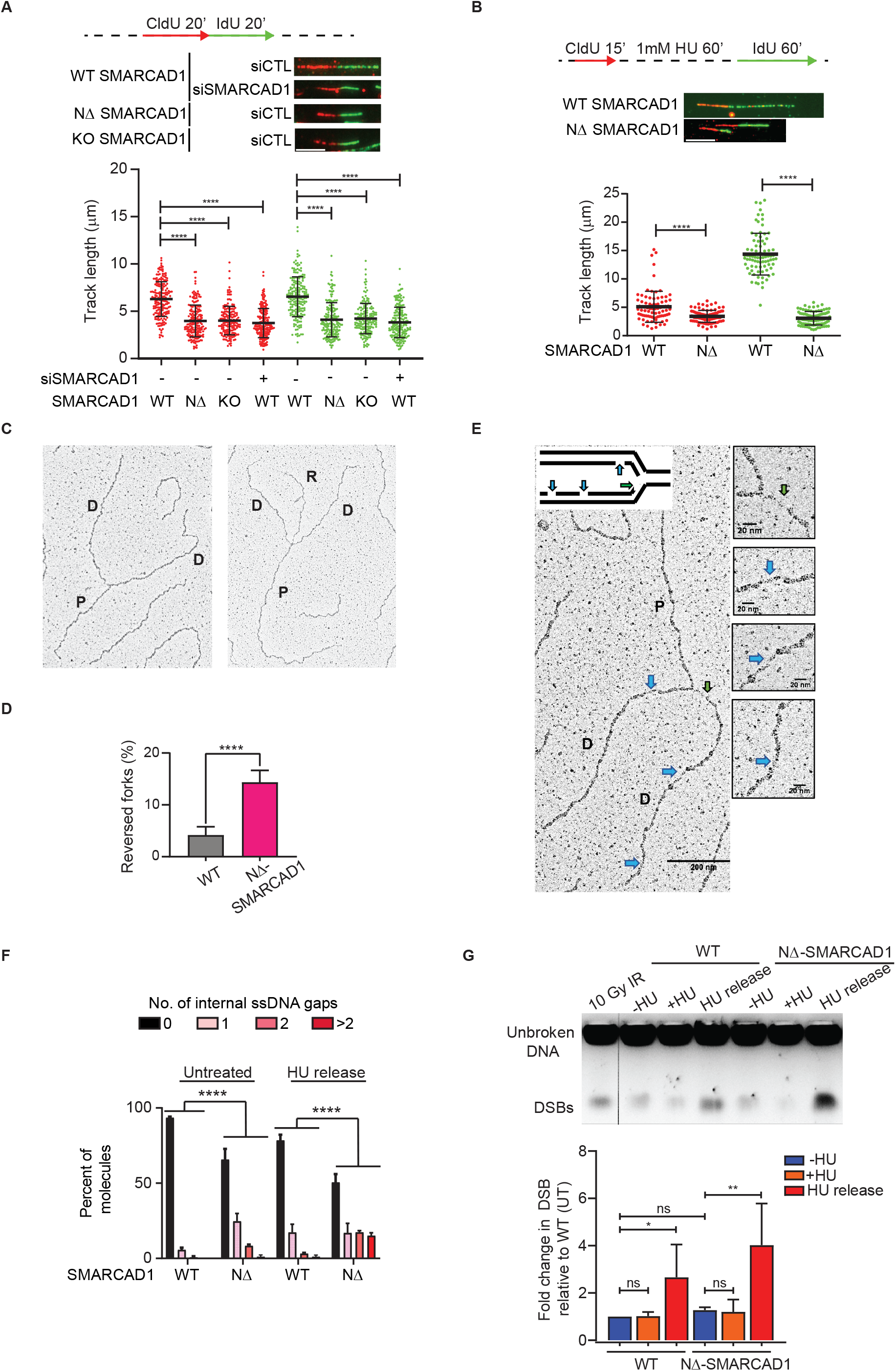
SMARCAD1 is required for proper fork progression, fork restart and genome stability. (**A**) Top panel: Schematic of replication fork progression assay with CldU and IdU labeling in WT, NΔ-SMARCAD1 and SMARCAD1^-/-^ (KO) cells. Representative DNA fibers for each condition are shown below the schematic (scale bar = 5 μm). Bottom panel: CldU (red) and IdU (green) track length (μm) distribution for the indicated conditions. (\*\*\*\**P* ≤ 0.0001, Kruskal-wallis followed with Dunn’s multiple comparison test, n= 3 independent experiment with similar outcomes). (**B**) Top panel: Schematic of replication fork restart assay. Representative DNA fibers for each condition are shown below the schematic (scale bar = 5 μm). Bottom panel: CldU (red) and IdU (green) track length (μm) distribution for the indicated conditions. (\*\*\*\**P* ≤ 0.0001, unpaired *t*-test). All DNA fiber experiments presented here were repeated three times with similar outcomes. (**C**) Representative image of a normal (left) and a reversed replication fork (right) observed by electron microscopy (EM). (D, daughter strand; P, parental strand; R, reversed arm). (**D**) Bar chart representing the percentage of fork reversal in WT and NΔ-SMARCAD1 cells in untreated condition. (\*\*\*\**P* ≤ 0.0001, unpaired *t*-test, n=3 independent experiments). (**E**) Representative electron micrographs of ssDNA gaps. (D, daughter strand; P, parental strand). Green and blue arrows point towards ssDNA gaps at the fork and behind the fork, respectively. (**F**) Bar chart representing the distribution of ssDNA gaps behind the fork in WT and NΔ-SMARCAD1 in untreated condition and 1hour after release from 1mM HU treatment. Chi-square test of trends was done to assess significance of internal ssDNA gaps between WT and NΔ-SMARCAD1 (****P < 0.0001, n=3 independent experiments). (**G**) Top panel: PFGE analysis for DSBs shows WT and NΔ-SMARCAD1 cells with and without 4mM HU treatment for 3 hours, and upon 16 hours release after the HU treatment. Bottom panel: Quantification from the three independent experiments showing DSB levels.

Since SMARCAD1 deficiency displayed significant replication defects during unperturbed replication (Fig. 3A and fig. S1I), we wondered if SMARCAD1 also plays a role in the progression after fork stalling. To assess the overall rate of DNA synthesis upon replication stress, we treated cells with 1mM HU for an hour. The replication rate after stress was measured by allowing the EdU incorporation for various time-points after release from HU and EdU intensities were measured in >3000 cells using high content microscopy. Upon 30 minutes of release from HU we observed a mild reduction in EdU incorporation in NΔ-SMARCAD1 cells. However, the reduction in EdU incorporation became more evident at later time points in NΔ-SMARCAD1 cells (fig. S2C). To further verify this, we performed a fork restart assay using DNA fiber analysis. Cells were labeled with CldU followed by a mild dose of HU (1mM) treatment for an hour to stall the forks and subsequently released into IdU. Consistently, we observed significant defects in CldU track lengths representing an internal control for unperturbed forks (Fig. 3B) similar to those observed in the fork progression assay performed in Fig. 3A. However, analysis of IdU track lengths representing stressed forks revealed an even higher shortening of the track lengths in NΔ-SMARCAD1 cells suggesting a more severe defect in the progression or restart of stalled forks (Fig. 3B). Additionally, upon analysis of fork restart efficiency, we observed a significant difference between stalled versus restarted forks in NΔ-SMARCAD1 cells (25% restarted) when compared to WT cells (60% restarted) after 15 minutes of release from HU-stress whereas this difference significantly reduced after 30 minutes of release from HU (86% WT, 74% NΔ-SMARCAD1) (fig. S2D, left) but the progression of restarted fork remained severely defective in NΔ-SMARCAD1 cells (fig. S2D, right). These data suggest that forks restart in absence of SMARCAD1 with moderate delay but further shows severe defects in progression of stressed forks. Thus, SMARCAD1 mediates both, the efficient restart as well as progression of replication forks, which also supports the finding that cells lacking SMARCAD1 are sensitive to replication stress inducing agents.

### SMARCAD1 prevents accumulation of under-replicated regions and consequent genome instability

To investigate whether the delayed restart and poor fork progression upon release from HU stress results in increased single-stranded DNA (ssDNA) levels in the NΔ-SMARCAD1 cells, we analyzed RPA32, a surrogate for ssDNA, by chromatin fractionation. Upon HU treatment, the RPA32 signals were markedly enhanced in WT cells (fig. S2E). Interestingly, untreated NΔ-SMARCAD1 cells showed a marked increase in chromatin associated RPA32 compared to untreated WT cells, suggesting that the accumulation of under-replicated regions in the genome could be due to defects in normal fork progression (Fig. 3A and fig. S2E). However, a significant increase in RPA32 levels could be seen upon HU treatment as well as upon release from HU-mediated block in NΔ-SMARCAD1 cells, suggesting that loss of SMARCAD1 at forks causes significant accumulation of under-replicated regions (fig. S2E).

DNA replication stress, exogenous or endogenous, results in reversal of forks (*32-35*), we hypothesized that slower fork progression and accumulation of RPA in NΔ-SMARCAD1 mutants in unperturbed conditions could be a result of frequent fork stalling that stabilizes into reversed forks. To test this hypothesis, we visualized replication intermediates formed *in vivo* using electron microscopy (EM) (*36*) in WT and NΔ-SMARCAD1 mutant cells. Interestingly, we observed a higher frequency of reversed forks in NΔ-SMARCAD1 than in WT cells, suggesting frequent stalling as well as remodeling of forks even in unperturbed conditions (Fig. 3, C and D). Moreover, we also observed an increase in the percentage of ssDNA gaps accumulated in daughter strands behind the fork of NΔ-SMARCAD1 cells relative to WT, which further enhanced dramatically upon release from HU mediated stress (Fig. 3E and F). We also quantified the length of ssDNA at the fork that determines nascent strand processing activity at the fork, which showed no significant difference in NΔ-SMARCAD1 than compared to WT (fig. S2F). Together, these data further corroborate that the role of SMARCAD1 is critical in limiting fork stalling under unperturbed conditions and promoting efficient fork restart as well as fork progression globally upon replication stress.

We further investigated whether the increased accumulation of ssDNA upon replication stress leads to an increase in DSBs that would contribute to genome instability. To evaluate the accumulation of DNA damage, we performed pulsed-field gel electrophoresis (PFGE) to measure the physical presence of DSBs. There was no obvious increase in the level of DSBs upon the stalling of forks induced by HU treatment in either WT and NΔ-SMARCAD1 cells, suggesting that forks stalled for 3-hours with HU treatment do not immediately collapse and convert into DSBs. This data was further supported by the efficient loading of RAD51 observed at stalled forks induced upon HU treatment in NΔ-SMARCAD1 similar to WT (fig. S2G). However, after release from replication stress for 16-hours, a marked increase in the signal of broken DNA fragments can be observed in NΔ-SMARCAD1 cells in comparison to WT cells (Fig. 3G). Together, these data suggest a role of SMARCAD1 at replication forks that is crucial to maintain genome integrity upon replicative stress.

### SMARCAD1 maintains PCNA levels at replication forks, especially upon fork restart

Since NΔ-SMARCAD1 lacks interaction with PCNA (Fig. 1F and fig. S1H) and NΔ-SMARCAD1 cells show defects in fork progression (Fig. 3, A and B), we wondered if the loss of SMARCAD1 at replication fork affects the PCNA clamp that acts as processivity factor for efficient DNA synthesis. We, therefore, measured the chromatin bound PCNA levels in replicating cells labelled with EdU to observe the dynamics of PCNA localization during DNA synthesis. QIBC analysis showed significant reduction in chromatin bound PCNA levels in replicating cells of NΔ-SMARCAD1 in comparison to WT (Fig. 4A), whereas the total levels of PCNA protein were not affected (Fig. 4B). This data suggests that absence of SMARCAD1 at forks affect PCNA levels at the forks. A similar reduction in PCNA levels at replication sites was observed in SMARCAD1^-/-^ cells suggesting NΔ-SMARCAD1 behaves similar to complete loss of SMARCAD1 protein and that NΔ-SMARCAD1 does not display a dominant negative phenotype (fig. S3A). We further monitored the impact of HU-mediated replication stress on PCNA recovery. Since PCNA dissociates from HU-mediated stalled forks (*8*) (Fig. 1, A and B), we hypothesized that aggravated defects in fork restart in NΔ-SMARCAD1 were due to poor recovery of PCNA at the forks upon release from HU. Using QIBC analysis, we simultaneously assessed the EdU incorporation and PCNA recovery upon HU stress using an average of 3000 cells per condition (Fig. 4C). WT replicating cells showed significantly reduced PCNA levels upon 1mM HU treatment for an hour and had recovered to their untreated levels by 45 minutes of release from HU stress (Fig. 4C and fig. S3B). Consistently, we observed reduced PCNA levels as well as reduced EdU incorporation in NΔ-SMARCAD1 cells in comparison to WT cells under the untreated condition. Interestingly, NΔ-SMARCAD1 cells showed severe defects in recovery of PCNA levels as well as reduced EdU incorporation upon release from HU-mediated replicative stress (Fig. 4C, and fig. S3, B and C). The significantly reduced EdU incorporation is consistent with the results of the DNA fiber assay of fork restart upon HU stress which revealed severe defects in the progression of restarted forks in NΔ-SMARCAD1 cells (Fig. 3B). This data suggests that SMARCAD1 participates in the maintenance of PCNA levels at the unperturbed forks. Moreover, under stressed conditions the absence of SMARCAD1 results in poor recovery of PCNA at restarting stalled forks, which subsequently causes inefficient fork restart and severe defects in fork progression upon replication stress.

**Fig. 4.**
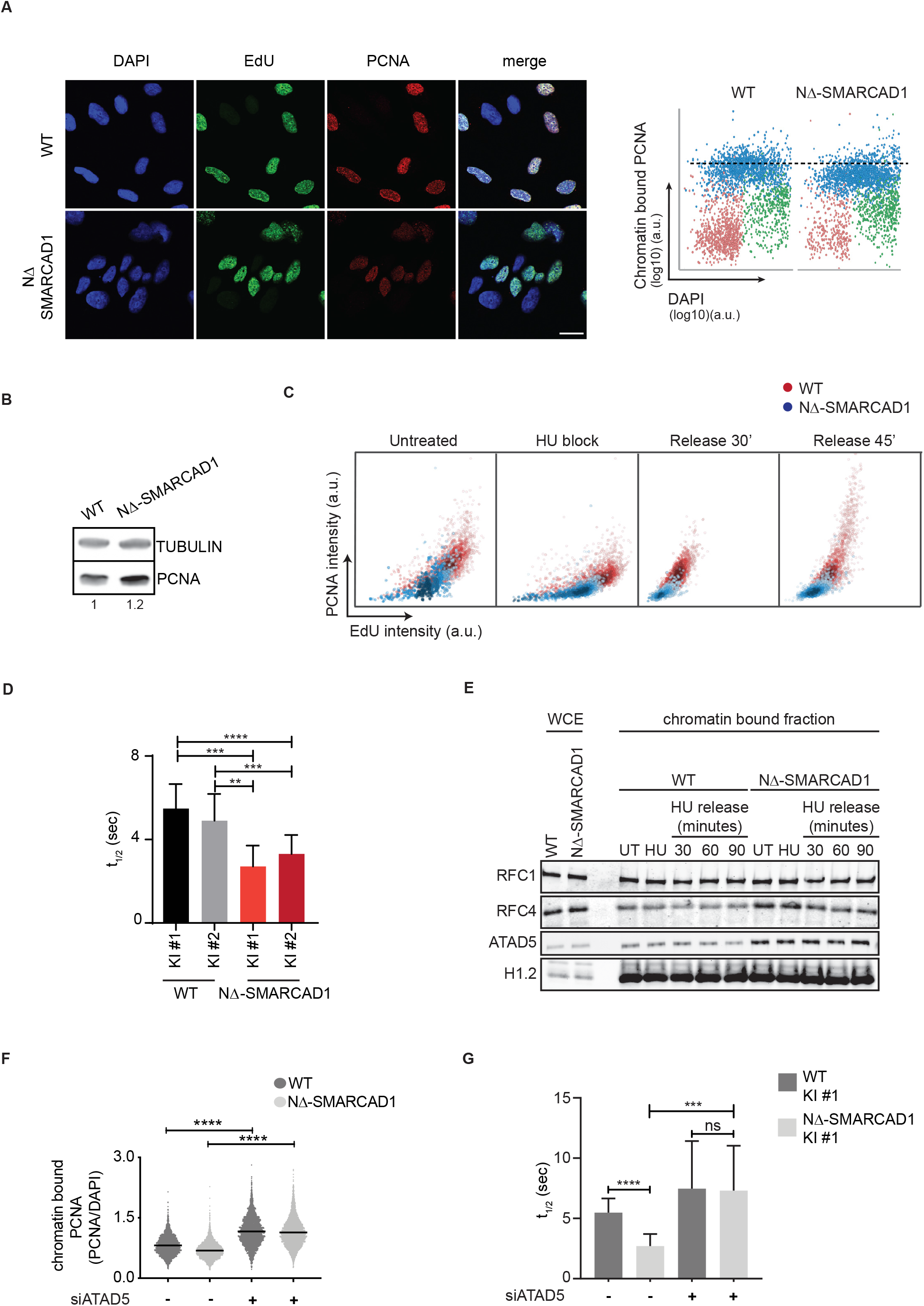
SMARCAD1 maintains PCNA level at replication forks. (**A**) Left: Representative confocal images showing chromatin bound PCNA (red) in EdU (green) positive WT and NΔ-SMARCAD1 MRC5 cells. Nucleus was stained with DAPI (blue) (scale bar = 20 μm). Right: QIBC analysis of the chromatin bound PCNA in WT and NΔ-SMARCAD1 cells. G_0-1_, S and G_2_-M phase cells are labeled in red, blue and green respectively. Dotted lines represent the mean chromatin bound PCNA intensity of S-phase cells in WT cells. (**B**) Immunoblot showing the total level of PCNA in WT and NΔ-SMARCAD1 cells. Tubulin is used as a loading control. Numbers below represent the quantification of PCNA level after normalized to the loading control. (**C**) QIBC analysis of PCNA vs EdU is shown in WT and NΔ-SMARCAD1 cells in untreated, 1mM 1hour HU block and 45 minutes release after HU conditions (note that for the HU block condition EdU labelling was performed prior to HU treatment). >1,800 S-phase cells were plotted in each condition. The color gradient represents the density of the cells. (**D**) Quantification of half-life of the GFP-PCNA fluorescence decay in GFP-tagged PCNA knock-in (KI) WT and NΔ-SMARCAD1 clones, mean±S.D. (\*\*\*\**P* ≤ 0.0001, \*\*\**P* ≤ 0.001, \*\**P* ≤ 0.01, unpaired *t*-test). (**E**) Immunoblot showing the whole cell extract (WCE) and chromatin bound fraction of RFC1, RFC4 and ATAD5 in WT and NΔ-SMARCAD1 cells. H1.2 is used as a loading control. (**F**) QIBC analysis of chromatin bound PCNA intensity (normalised to DAPI) in si-control and si-ATAD5 treated WT and NΔ-SMARCAD1 cells. (\*\*\*\**P* ≤ 0.0001, unpaired *t*-test). (**G**) Quantification of half-life of the GFP-PCNA fluorescence decay in GFP-tagged PCNA KI WT #1 and NΔ-SMARCAD1 #1 cells treated with or without siATAD5. Mean±S.D. (\*\*\*\**P* ≤ 0.0001, \*\*\**P* ≤ 0.001, unpaired *t*-test).

We further determined the dynamics of PCNA in replicating WT and NΔ-SMARCAD1 cells using an inverse Fluorescence Recovery After Photobleaching (iFRAP) live-cell imaging assay. iFRAP is an adapted FRAP approach optimized to analyze differences of dissociation rates (K_off_) and involves continuous bleaching to quench the total nuclear fluorescence of a GFP-tagged protein with the exception of a small predefined area. Using this approach, we could determine the residence time of GFP-PCNA at replication foci (unbleached area) as a direct read out of its turnover (fig. S3D). We performed iFRAP on GFP-tagged PCNA expressed from its endogenous allele in both WT and NΔ-SMARCAD1, cell types (fig. S1G). Remarkably, we observed nearly 2-fold shorter residences times for GFP-tagged PCNA foci in NΔ-SMARCAD1 cells compared to WT cells (Fig. 4D and fig. S3D). This data clearly suggests that the turnover of PCNA at replication forks is severely increased in the absence of SMARCAD1 at the forks, which may be caused by either a defect in the loading or unloading of PCNA in the absence of SMARCAD1 at the replication forks.

To further test this hypothesis, we performed chromatin fractionation to observe the chromatin-associated fraction of subunits of the PCNA loader, RFC (RFC1/RFC2-5) and of the unloader, RLC (ATAD5/RFC2-5) complex subunits (*15, 37*). We observed no obvious change in the level of RFC1, a major subunit of the RFC complex, in either cell type with or without HU treatment (Fig. 4E). Interestingly, the chromatin association of RFC4, a subunit shared between the RFC and RLC complexes, as well as that of ATAD5, a major subunit of the RLC complex, were found to be significantly enhanced in chromatin bound fraction of NΔ-SMARCAD1 cells, while the total level of these proteins in whole cell extracts remains similar to WT (Fig. 4E). This finding suggests that the increased chromatin binding of the PCNA-unloader ATAD5-RLC causes the increased release of PCNA in the absence of SMARCAD1. To further rule out the possibility of deregulated mRNA expression of ATAD5-RLC complex in SMARCAD1 mutants, we compared the transcriptome analysis data showing similar level of PCNA, ATAD5, RFC1 and all the other RFC subunits (RFC2-5) that are shared between loading and unloading complexes (fig. S3E). Based on this observation, we next tested whether depleting ATAD5 levels might restore normal PCNA chromatin association and reduce replication defects in NΔ-SMARCAD1 cells. Consistent with previous reports (*38*), we observed enhanced PCNA levels at replicating sites in WT cells upon ATAD5 knockdown using high content microscopy (Fig. 4F and fig. S3F). Importantly, ATAD5 knockdown rescued PCNA levels at replication sites in NΔ-SMARCAD1 cells, similar to WT levels (Fig. 4F and fig. S3F). We further confirmed these observations using iFRAP and detected an increased retention time of PCNA in both WT and NΔ-SMARCAD1 cells (Fig. 4G). However, as previously reported (*38*), loss of ATAD5 significantly reduced the overall EdU incorporation in WT cells and a similar decrease was observed in NΔ-SMARCAD1 cells, suggesting that the enhanced accumulation of PCNA at forks also affects overall DNA synthesis (fig. S3G). Furthermore, the ATAD5 knockdown did not rescue the cellular sensitivity of NΔ-SMARCAD1 cells to cisplatin and olaparib (fig. S3H).

### Loss of 53BP1 restores PCNA stability, fork restart and drug tolerance in NΔ-SMARCAD1 cells

Having established the role of SMARCAD1 at the replication forks, we further investigated the mechanism of how SMARCAD1 promotes replication fork progression. Earlier studies have shown a role for SMARCAD1 in displacing 53BP1 from the site of DSBs to promote HR repair (*23*). Moreover, SMARCAD1 and 53BP1 show contrasting enrichments at unperturbed versus stalled replication forks, shown by iPOND-SILAC-Mass Spectrometry (*8*) (Fig. 1A and Table S1). We further validated the enrichments of 53BP1 at stalled forks versus restarted forks using fluorescence microscopy in WT cells (Fig. 5A). The data clearly showed 53BP1 colocalization with EdU mainly upon HU treatment suggesting its enrichment at stalled forks in WT cells, whereas upon release from HU stress, the EdU labelled sites representing restarted forks show clear displacement between 53BP1 and EdU foci (Fig. 5A). We hypothesized that, similar to DSBs (*23*), SMARCAD1 might prevent 53BP1 to accumulate at active or restarted replication forks by promoting its displacement from the stalled forks. To test this hypothesis, we measured the levels of 53BP1 protein in replicating cells (EdU positive) of NΔ-SMARCAD1 compared to WT, in untreated as well as in cells released from HU-stress. We observed a mild but significant increase in 53BP1 levels in replicating cells of NΔ-SMARCAD1 and strikingly, a significantly higher accumulation of 53BP1 levels could be seen in cells released from HU-stress (fig. S4A). We further measured the localization of 53BP1 protein relative to EdU marked replication sites in NΔ-SMARCAD1 compared to WT cells. Upon HU block, a significant percentage of replicating WT cells showed an overlap between EdU and 53BP1 foci, which significantly reduced upon release from HU stress (Fig. 5B). Whereas significantly higher percentage of NΔ-SMARCAD1 cells showed colocalization of EdU and 53BP1 foci in HU block cells, which remained remarkably higher even upon release from HU stress (Fig. 5B). Supporting this observation, the Pearson’s overlap co-efficient as well as Manders’ (M1/M2) overlap co-efficients estimating the significance of overlap between EdU and 53BP1 foci were found to be significantly higher in NΔ-SMARCAD1 than in WT (fig. S4B). Together these data suggest that SMARCAD1 is required to displace 53BP1 from stalled replication forks possibly to allow their restart.

**Fig. 5.**
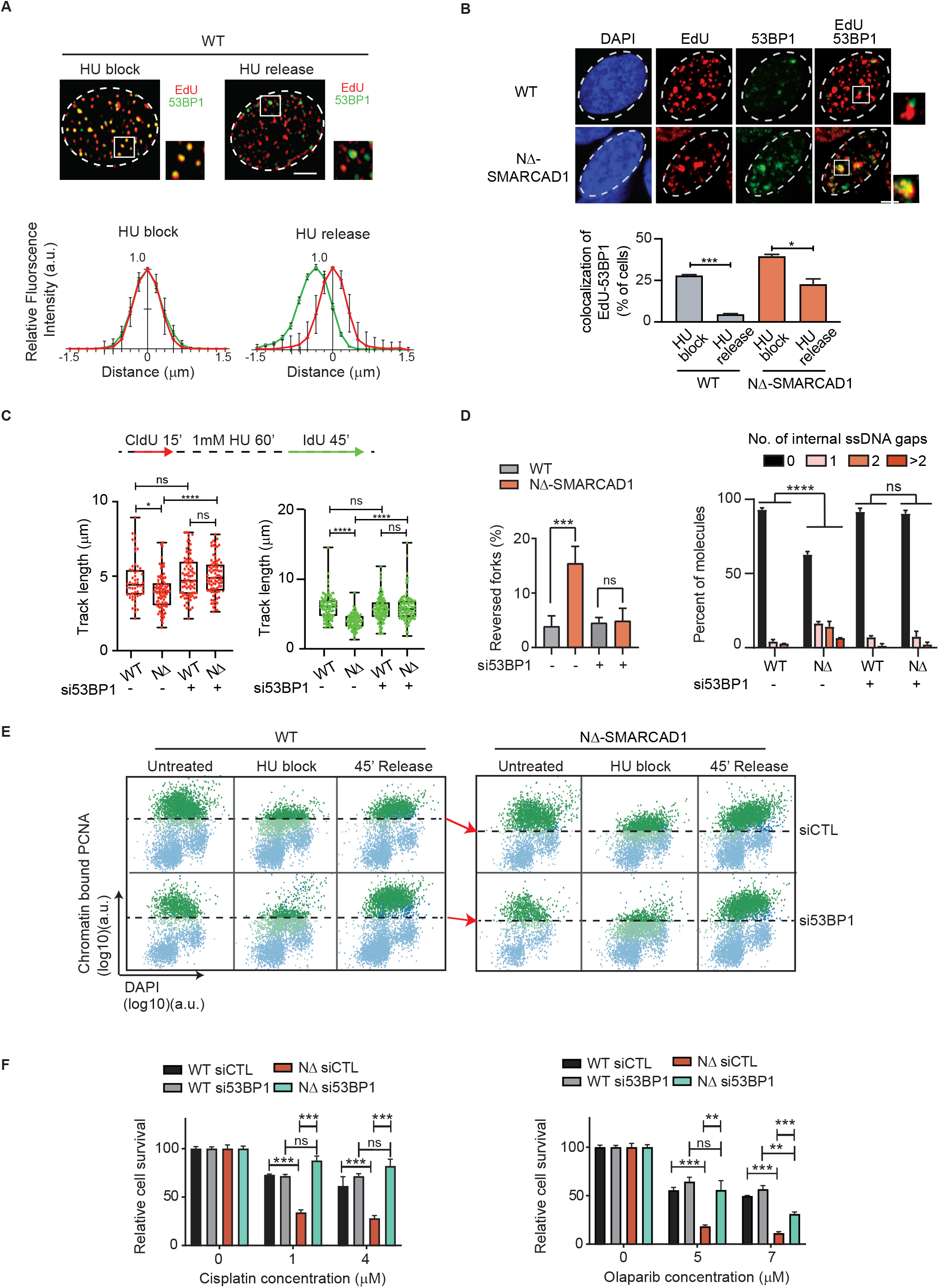
SMARCAD1 prevents 53BP1 enrichment at forks to maintain PCNA levels, fork progression and genome stability. (**A**) Top Panel: Representative image showing 53BP1 (green) and EdU (red) in WT cells treated with 4mM HU for 3hour (HU block) and 1 hour after release from HU block (HU release) (scale bar = 5 μm). Bottom Panel: The average distance between EdU and 53BP1 foci in HU block and HU release condition is shown. Error bars stand for ±S.D. (**B**) Top Panel: Representative confocal images showing DAPI (blue), EdU (red) and 53BP1 (green) in HU release condition in WT and NΔ-SMARCAD1 cells (scale bar = 5 μm). Bottom Panel: Quantification of cells showing EdU and 53BP1 co-localisation in WT and NΔ-SMARCAD1 cells with 4mM 3hour HU block and with 1-hour HU release condition. (\*\*\**P* ≤ 0.001,\**P* ≤ 0.05, unpaired *t*-test). (**C**) Top panel: Schematic of replication fork restart assay. Bottom panel: CldU (red) and IdU (green) track length (μm) distribution for the indicated conditions. (\*\*\*\**P* ≤ 0.0001, \**P* ≤ 0.05,, ns, non-significant, Kruskal-wallis followed with Dunn’s multiple comparison test, n= 3 independent experiment with similar outcomes). (**D**) Left: The frequency of reversed forks was quantified using electron microscopy in WT and NΔ-SMARCAD1 cells with or without 53BP1 knock down. (\*\*\*\**P* ≤ 0.0001, ns, non-significant, unpaired *t*-test). Right: Bar chart representing the distribution of ssDNA gaps behind the fork of si-control or si-53BP1 treated WT and NΔ-SMARCAD1 cells. (\*\*\*\**P* ≤ 0.0001, ns, non-significant, Chi-square test, n= 3 independent experiment). (**E**) QIBC analysis of chromatin bound PCNA dynamics and DAPI in untreated, HU block and HU release condition of si-control and si-53BP1 in WT and NΔ-SMARCAD1 cells. Cells above dotted lines represent the PCNA positive S-phase cells in WT and NΔ-SMARCAD1 cells. The red arrows compare the level of PCNA in WT and NΔ-SMARCAD1 cells upon si-control and si-53BP1 conditions. (Note that for the HU block condition EdU labelling was performed prior to HU treatment) (**F**) Quantification of colony survival assay of si-control and si-53BP1 in WT, NΔ-SMARCAD1 cells treated with different concentrations of olaparib and cisplatin. Error bars stand for + S.D. (n=3). (\*\*\**P* ≤ 0.001, \*\**P* ≤ 0.01, ns, non-significant, unpaired *t*-test).

This observation led us to hypothesize that loss of 53BP1 may allow the normal progression of forks in NΔ-SMARCAD1 cells, which shows frequent fork stalling even in unperturbed conditions (Fig. 3C). We, therefore, first investigated the progression rate of unperturbed forks using si53BP1 in NΔ-SMARCAD1 using a DNA fiber assay. Interestingly, transient knockdown of 53BP1 completely rescued the fork progression defects of NΔ-SMARCAD1 cells (fig. S4, C and D). Additionally, we also performed fork restart assay and found that both the IdU track lengths as well as CldU track lengths, representing stressed (after HU treatment) and non-stressed forks (before HU treatment) respectively, showed complete restoration of fork progression rates in NΔ-SMARCAD1 (Fig. 5C). Consistently, we observed a rescue in accumulation of reversed forks as well as reduced accumulation of ssDNA gaps behind the fork in NΔ-SMARCAD1 cells upon 53BP1 knock down condition (Fig. 5D). As the severe defects in restart of replication forks in NΔ-SMARCAD1 was correlated with the poor recovery of PCNA, we next sought to determine, if 53BP1 knockdown would also restore PCNA levels in NΔ-SMARCAD1 cells. Consistently, QIBC plots showed that upon HU-mediated block PCNA levels were significantly reduced in replicating cells even upon 53BP1 knockdown, however, importantly, QIBC plots showed a remarkable recovery of PCNA in NΔ-SMARCAD1 similar to WT, when released from HU-mediated block (Fig. 5E and fig. S4E). In support to the restoration of PCNA levels, we observed a marked reduction in chromatin bound ATAD5 levels upon knockdown of 53BP1 in NΔ-SMARCAD1 (fig. S4F), suggesting that 53BP1 further promotes PCNA unloading in absence of SMARCAD1 at forks through ATAD5 activity. The potential interaction between 53BP1 and ATAD5 was further confirmed by chromatin immunoprecipitation of 53BP1 showing enhanced interaction in either HU-induced replication stress conditions in WT or under unperturbed conditions of NΔ-SMARCAD1 cells, both of which shows enhanced accumulation of stalled forks (Fig. 3C and fig. S4G). We also noticed that the higher molecular weight band of ATAD5 was mainly immunoprecipitated with 53BP1 in chromatin IPs which was further confirmed by notable reduction in signal of potentially phosphorylated ATAD5 band in cells targeted with siATAD5 (fig. S4G). The phosphorylated form of ATAD5 have been reported to interact with RAD51 at stalled/regressed forks previously (*39, 40*). Taken together, these data suggest that 53BP1 interaction with ATAD5 regulates PCNA levels at stalled forks. Since loss of 53BP1 rescued genome instability, as monitored by reduction of accumulated ssDNA gaps in NΔ-SMARCAD1 (Fig. 5D), we next determined if 53BP1 knockdown rescues the sensitivity of NΔ-SMARCAD1 cells towards replication poisons. Interestingly, we observed a significant restoration of resistance towards cisplatin and olaparib treatment after depletion of 53BP1 in NΔ-SMARCAD1 cells (Fig. 5F). Together, these data imply that SMARCAD1 maintains fine PCNA levels by suppressing unscheduled 53BP1 accumulation at the active replication forks and thereby maintain genome stability and replication stress tolerance in the cells.

From this data, we further hypothesized that enzymatic activity of SMARCAD1 is required to displace 53BP1-associated nucleosomes to suppress the accumulation of 53BP1 at replication forks, in order to promote efficient fork restart and progression. To investigate this, we generated knock-Ins of cDNA-SMARCAD1 that were either wildtype or contained an ATPase-disabling K528R mutation (*23*). As expected, we observed a rescue in fork progression defects in NΔ-SMARCAD1 cells when corrected with fully functional SMARCAD1 but not with ATPase-dead K528R SMARCAD1 (fig. S4H). Moreover, ATPase-dead SMARCAD1 showed significant defects in fork progression when it replaced wildtype SMARCAD1 in WT cells (fig. S4H), suggesting that the ATPase chromatin remodeling activity of SMARCAD1 is essential to maintain fork stability.

### SMARCAD1-mediated active fork stability confers survival in BRCA1 mutated tumors, irrespective to their HR-status

Our data implies that SMARCAD1-mediated replication fork stability contributes to genome stability in a manner independent of its role in HR repair of DSBs. Similarly, HR-independent roles in the protection of stalled forks during replication stress have been uncovered for BRCA1 and BRCA2 (*2, 3, 5-7*). To further test if SMARCAD1 also protects stalled forks, similar to BRCA1, we observed for fork degradation using DNA fiber assay. The data clearly shows that loss of BRCA1 leads to stalled fork degradation even upon 3h exposure to 4mM HU, while NΔ-SMARCAD1 shows no significant defects in fork protection and is similar to WT (Fig. 6A). Furthermore, as shown previously longer exposure of cells to 4mM HU (up to 8hr) leads to a moderate but significant processing of forks in WT cells (*41*), we observed similar effects in NΔ-SMARCAD1 while loss of BRCA1 led to severe fork degradation (Fig. 6A). Further, this data also suggests that SMARCAD1 is not defective in processing of stalled forks, as proposed for its fission yeast homolog (*42*), otherwise the moderate but significant degradation observed in 8hours similar to WT level would not be expected due to defective processing of nascent strands. Thus, these data along with fork progression data (Fig. 3, A and B) taken together suggest that replication defects observed in absence of SMARCAD1 is due to defective active replication fork stability and not due to defective stalled fork protection or fork processing activities. Furthermore, in the absence of SMARCAD1, unperturbed cells showed frequent stalling of replication forks without subsequent accumulation of DSBs (Fig. 3, C and G), this could possibly be due to BRCA-mediated fork protection in SMARCAD1 mutant cells. To test this hypothesis, we knocked down BRCA1 transiently from MRC5 WT, NΔ-SMARCAD1 and SMARCAD1^-/-^ cells to analyze replication fork dynamics. As previously reported, siBRCA1 in WT cells showed no significant defects in progression rate of unperturbed forks (*2*). However, in NΔ-SMARCAD1 and SMARCAD1^-/-^ cells, loss of BRCA1 resulted in significantly shorter track length (fig. S5A), which could not be rescued by loss of 53BP1 (fig. S5B). These data suggest that upon loss of SMARCAD1, BRCA1 is required to maintain progression of forks, possibly by protecting stalled forks from DNA nuclease mediated degradation to allow their restart. To test if indeed loss of BRCA1 in SMARCAD1 mutants lead to increased DNA damage, we performed QIBC analysis for γH2AX, and observed a significantly enhanced accumulation of DNA damage upon BRCA1 knockdown in both NΔ-SMARCAD1 as well as SMARCAD1^-/-^ mutants compared to single mutants or wildtype cells (Fig. 6B), suggesting BRCA1 could be required to protect stalled forks from degradation to prevent DNA damage accumulation.

**Fig. 6.**
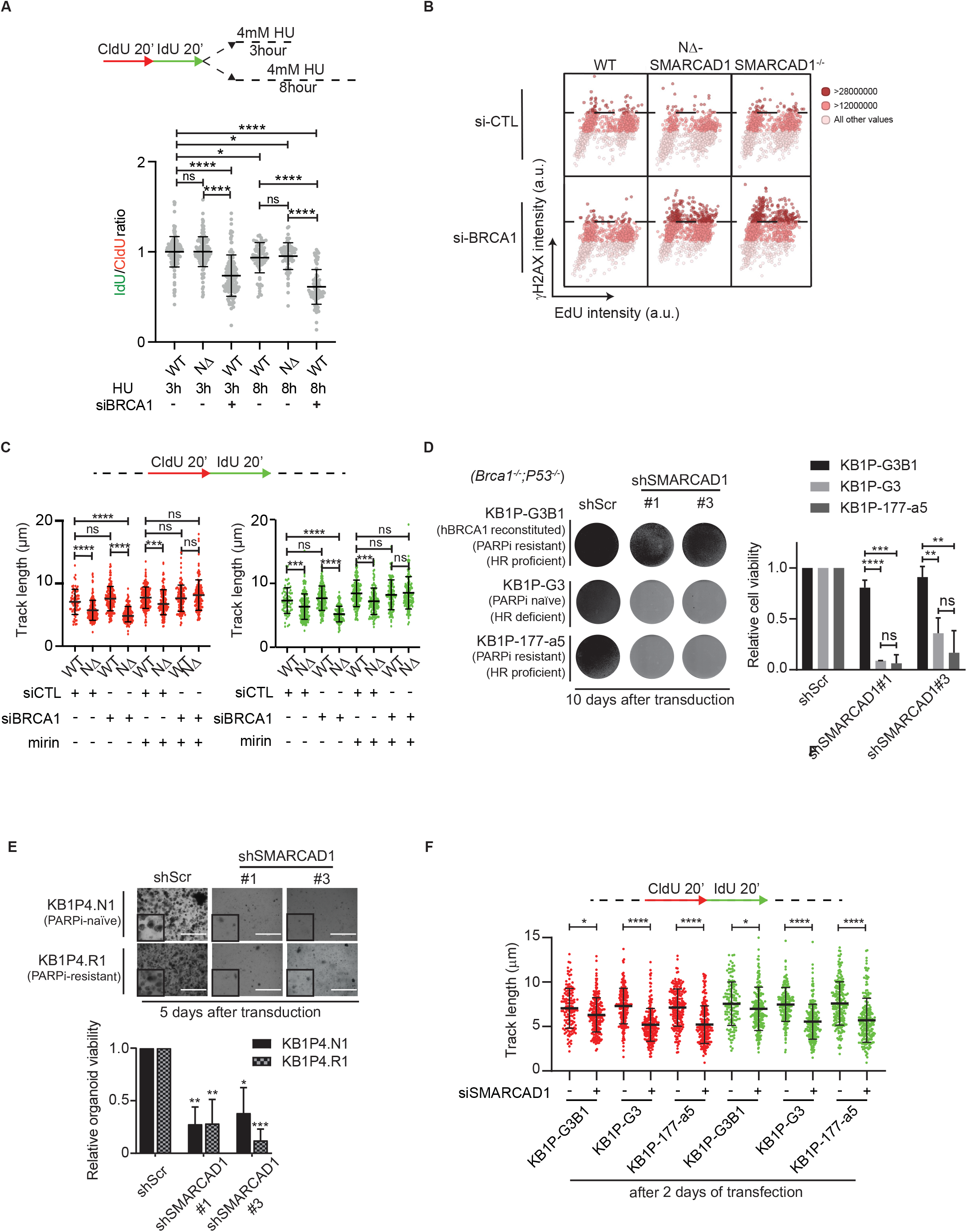
Smarcad1 is essential for fork progression and proliferation of BRCA1 deficient mouse tumor cells. (**A**) Top panel: Schematic of replication fork degradation assay with CldU and IdU labeling. Bottom panel: Ratio of IdU to CldU tract length was plotted for the indicated conditions. (**B**) QIBC analysis of γH2AXvs EdU is shown in WT, NΔ-SMARCAD1 and SMARCAD1^-/-^ cells in si-control and si-BRCA1 conditions. >1,000 cells were plotted in each condition. The color gradient represents the γH2AX levels in each cells. (**C**) Top panel: Schematic of replication fork progression assay with CldU and IdU labeling. Bottom panel: CldU (red) and IdU (green) track length (μm) distribution for the indicated conditions. (\*\*\*\**P* ≤ 0.0001, \*\*\**P* ≤ 0.001, \**P* ≤ 0.05, ns, non-significant Kruskal-wallis followed with Dunn’s multiple comparison test, n= 3 independent experiment with similar outcomes). (**D**) Left: Representative images of KB1P (*Brca1*^*-/-*^ *P53*^*-/-*^) mouse tumor cells pooled from three independent experiments at day 3 and imaged at day 10, after transduction of scramble control shRNA and shSMARCAD1#1 and #3. Right: Quantification of cell viability using crystal violet staining assay. Error bars stand for + S.D. (n=3). (\*\*\*\**P* ≤ 0.0001, \*\*\**P* ≤ 0.001, \*\**P* ≤ 0.01, ns, non-significant, unpaired *t*-test). (**E**) Top panel: Representative images of KB1P mouse tumor organoid. Image taken 5 days after the transduction of scramble control shRNA and shSMARCAD1#1 and #3 (scale bar = 1000μm). Bottom panel: Quantification of cell viability using cell titer blue assay. Error bars stand for + S.D. (n=3). (\*\*\**P* ≤ 0.001, \*\**P* ≤ 0.01, \**P* ≤ 0.05, unpaired *t*-test). (**F**) Top panel: Schematic of replication fork progression assay. Bottom panel: CldU (red) and IdU (green) track length (μm) distribution in KB1P mouse tumor cells treated with si-control or si-SMARCAD1. (\*\*\*\**P* ≤ 0.0001, \**P* ≤ 0.05, Kruskal-wallis followed with Dunn’s multiple comparison test, n= 3 independent experiment with similar outcomes).

As previously reported, BRCA1 protects stalled forks from degradation mediated by DNA nuclease Mre11 (*7*). Therefore, to test out this hypothesis, we treated cells with inhibitor of DNA nuclease Mre11, Mirin and monitored the fork progression using DNA fiber assay. Strikingly, Mirin treatment completely rescues the severe fork progression defects observed upon loss of BRCA1 in SMARCAD1 mutant (Fig. 6C). This data suggests that indeed stalled forks in absence of SMARCAD1 required BRCA1 protection to allow fork progression and maintain genome integrity.

Previously, SMARCAD1 was reported to play a critical role in the metastasis of triple-negative breast cancer (*43, 44*). To test whether differential levels of SMARCAD1 expression could be an indicator of patient responses to replication stress inducing platinum chemotherapy, we analyzed the high grade serous ovarian cancer (HGSOC) patients for their correlation between BRCA1 and SMARCAD1 expression levels to their response to chemotherapy. Interestingly, survival analysis demonstrated that platinum-treated BRCA1-low patients, but not BRCA1-high patients, with low SMARCAD1 expression were correlated with a longer progression-free survival (PFS) while higher expression of SMARCAD1 correlated with poor response to chemotherapy (fig. S5C). This data suggests that SMARCAD1 levels could be a biomarker for acquired resistance to platinum-based chemotherapy in BRCA1-low/deficient ovarian cancers.

To further verify this experimentally, we queried if SMARCAD1 is required for fork progression in BRCA1-deficient tumor cells and whether its loss could hypersensitize HR-deficient BRCA1^-/-^ mouse breast tumor cells generated using *K14Cre;Brca1*^*F/F*^; *p53*^*F/F*^ (KB1P) mouse mammary tumor models (*45*). We generated shRNA-mediated knockdowns of SMARCAD1 in *Brca1*^*-/-*^ *P53*^*-/-*^ defective mouse breast tumor derived cell lines (fig. S5D). Surprisingly, the loss of SMARCAD1 resulted in a significant reduction in colony formation in HR-defective BRCA1^-/-^ (KB1P-G3; PARPi naïve) (*46*) tumor cells but not in KB1P-G3 tumor cells that were reconstituted with human BRCA1 (KB1P-G3-B1) and proficient in HR (*47*), suggesting that loss of SMARCAD1 causes synthetic lethality in BRCA1-deficient tumor cells (Fig. 6D). These data indicate a potential role of SMARCAD1 in maintaining active fork stability, which may be the reason for the survival of BRCA1-deficient HR-defective tumor cells. Furthermore, we also tested whether BRCA1 and 53BP1 double-knockout tumor cells which are proficient for HR and resistant to PARPi treatments (KB1P-177.a5; PARPi resistant) (*46*), require SMARCAD1 for proliferation. Interestingly, a SMARCAD1 knockdown again resulted in lethality in these cells, suggesting that SMARCAD1’s role is essential for proliferation of BRCA defective tumor cells, irrespective of their HR status (Fig. 6D). Furthermore, 53BP1 deficiency in BRCA1-defective genetic background could not rescue defects of SMARCAD1 knockdown, which suggests that fork protection mediated by BRCA1 becomes critical for cellular survival in the absence of SMARCAD1, similar to what we observed in human fibroblast cells (fig. S5, A and B). Additionally, we tested the effect of SMARCAD1 knockdown on KB1P -derived, PARPi-naïve (KB1P4.N) and PARPi-resistant (KB1P4.R), tumor organoids grown in ex vivo cultures (*48*). Consistent with our results in KB1P tumor cell lines, we observed a synthetic lethality in the 3D-tumor organoids, suggesting that SMARCAD1 is essential for the survival of BRCA1-mutated tumors (Fig. 6E). These data strongly suggest a conserved and non-epistatic role of SMARCAD1 and BRCA1 at replication forks.

BRCA1-deficient cells show reduced fork protection and high levels of endogenous stress (*7, 49*), we speculated that the loss of SMARCAD1 further enhances replication stress due to defective progression of forks causing proliferation defects. To test this speculation, we used siRNA to transiently deplete SMARCAD1 protein (*50*) in KB1P 2D-tumor derived cell lines (fig. S5E) to monitor individual fork progression using DNA fiber assay. We sequentially labeled human BRCA1-reconstituted, KB1P-G3B1 cells as control, KB1P-G3 (HR deficient) and KB1P-177.a5 (chemoresistant; HR proficient) with CldU (red) and IdU (green), followed by track length analysis. In support to the survival assays, even though sub-lethal SMARCAD1 knock-down affects only mildly the cell cycle of all 3 cell lines (fig. S5F), it led to a significantly shorter track lengths of both CldU and IdU in both KB1P-G3 and KB1P-177 cells in comparison to BRCA1 reconstituted KB1P-G3B1 cells, suggesting an essential role of SMARCAD1 in mediating fork progression in absence of BRCA1 (Fig. 6F). Together, these results strongly suggest that the SMARCAD1-mediated stability of active replication forks is a physiologically important process for cellular proliferation of BRCA1-deficient tumors, irrespective of their HR-status (fig. S6).

## DISCUSSION

Our study has revealed a novel mechanism of active fork stability that has important implications in the survival of tumor cells.

### A genetically distinct role of SMARCAD1 at active replication forks, from HR

As opposed to the commonly attributed role of DNA repair factors in replication fork protection (*6, 7, 9, 51*), here we show a newly recognized function of SMARCAD1 in maintaining the stability of active (unperturbed and restarted) replication forks while its absence do not disturb stalled fork protection and fork processing activities (Fig. 3, A-B and Fig. 6A and fig. S2F). Importantly, using a separation-of-function SMARCAD1 mutant (NΔ-SMARCAD1), we show that SMARCAD1’s role in stabilization of active replication forks is genetically separable from its role in HR repair, and is critical in maintaining genome stability especially upon replication stress. The physical interaction between SMARCAD1 and PCNA, established using in vitro and in vivo assays (*24*), was suggested to be responsible for SMARCAD1’s association with replication machinery (*24, 26*). Our biochemical and immunofluorescence assays further confirm that the NΔ-SMARCAD1 protein, lacking initial 137 amino acids, can bind to chromatin, but lacks the ability to interact with PCNA. This finding is consistent with the lack of association between NΔ-SMARCAD1 and replication forks as previously suggested (*26*). However, other components may also be involved in promoting SMARCAD1’s association with replication machinery, such as phosphorylation of SMARCAD1 by Cyclin-dependent kinase (CDK). Indeed a CDK phosphorylation site at the N-terminus of SMARCAD1 is among the 137 amino acids that are missing in the NΔ-SMARCAD1 protein (*52*). Nonetheless, the CUE-dependent protein-protein interactions and ATPase-dependent chromatin remodeling activity, in the context of HR repair and nuclear association, seems to remain functional in the NΔ-SMARCAD1 protein. Notably, cells with a SMARCAD1-null (SMARCAD1^-/-^) genotype and those expressing the NΔ-SMARCAD1 allele, show similar defects in fork progression and in sensitivity towards replication poisons, arguing that the role of SMARCAD1 at replication forks is crucial in mediating resistance to replication stress-inducing drugs.

Furthermore, our data showed evidence of frequent accumulation of stalled forks as well as ssDNA gaps behind the replication forks in NΔ-SMARCAD1 cells. The accumulation of ssDNA and stalled forks could be indicative of hindered replication fork progression through certain difficult-to-replicate regions, such as highly transcribing regions or repetitive regions of the genome (*53*). Alternatively, ssDNA accumulation could also be resultant of the re-priming events by PRIMPOL at stalled forks that in the process of re-initiating the DNA synthesis leads to accumulation of ssDNA gaps (*54, 55*). Interestingly, however in BRCA1-challenged cells, PRIMPOL activity was shown to be responsible for DNA synthesis upon replicative stress condition. Here, our study shows a unique pathway of active fork stabilization mediated by SMARCAD1 which is critical for fork progression in BRCA1-deficient cells even under unperturbed conditions. This implies that SMARCAD1 mediated active replication fork stability is a central and a separate pathway for stabilization of replication forks than from recently described PRIMPOL mediated fork re-priming or well-established BRCA1-mediated fork protection pathway (*56*).

### SMARCAD1 regulates PCNA levels at active replication forks

Our findings suggest a hitherto unrecognized role for SMARCAD1 in maintaining the fine control of PCNA levels at the forks. In this study, along with previously published study (*24, 26*), we have strong evidence of positive interaction between SMARCAD1 and PCNA which is also responsible for SMARCAD1 association with replication machinery. A global reduction in chromatin bound PCNA levels at the fork and a faster dissociation rate of PCNA foci in NΔ-SMARCAD1 cells, further suggests a mutualistic interaction between SMARCAD1 and PCNA at the replication forks (Fig.4, C-D). Consistently, an increase of PCNA unloading by the ATAD5-RLC complex was observed in NΔ-SMARCAD1 cells. A recent report demonstrated a critical role of ATAD5 in the removal of PCNA from stalled forks to promote recruitment of fork protection factors (*39*). Consistent with this report, we observed reduced PCNA levels at replication forks, accompanied by an increased accumulation of ATAD5-RLC complex, and increased frequency of reversed forks (protected stalled forks) in unperturbed NΔ-SMARCAD1 cells. Furthermore, a significant number of peptides arising from RFC2-5 protein subunits that are shared between PCNA loading (RFC) and unloading (ATAD5-RLC) complexes, were obtained from SMARCAD1 co-immunopurification (*24*). This data may indicate the direct involvement of SMARCAD1 in regulating loading/unloading activity of PCNA at replication forks. However, an interesting finding from our study is that loss of 53BP1 results in a significant restoration of PCNA levels in NΔ-SMARCAD1 cells accompanied with a significant reduction in ATAD5 levels at replication forks. Furthermore, the enhanced interaction between 53BP1 and post-translationally modified ATAD5 in HU treated wildtype cells or in unperturbed NΔ-SMARCAD1 cells seems to be regulating PCNA unloading from the forks. Whether the post-translation modification of ATAD5 are solely ATR-mediated or additional mechanisms play role in its regulation as suggested previously (*39*) could distinguish between the physiological role of ATAD5 in regulating PCNA dynamics that involves continuous loading/unloading events during normal fork progression versus the persistent unloading of PCNA from stalled forks.

### SMARCAD1 prevents 53BP1 accumulation to mediate tolerance to replication stress

Our study shows an unforeseen role of SMARCAD1 in preventing 53BP1 accumulation at active restarted replication forks. SMARCAD1 has been shown to displace 53BP1 from DSBs possibly by the displacing of the H2A-Ub nucleosomes with which 53BP1 associates (*23*). This observation is consistent with the finding that SMARCAD1 homologs in yeast perform nucleosome sliding and promote H2A-H2B dimer exchange in vitro, also regulating histone turnover in replicating cells of fission yeast cells (*57-59*). Consistent with these observations, it has been shown that the loss of SMARCAD1 results in a prolonged enrichment of 53BP1 at DSBs (*23, 29*). Strikingly, we found increased 53BP1 in association with restarted forks in NΔ-SMARCAD1 cells. Intriguingly, SMARCAD1 and 53BP1 also show contrasting enrichments at stalled versus unperturbed forks suggesting that their co-existence is possibly also prohibited by remodeling activity of SMARCAD1 at replication forks in a manner similar to that of their interaction at DSBs (*8*) (Fig. 1, A, C and Fig. 5A). Consistently, the knockdown of 53BP1 or the introduction of fully-functional SMARCAD1 but not the ATPase-dead SMARCAD1, results in the resumption of normal progression rates in NΔ-SMARCAD1 cells. This data implies that both the ability of SMARCAD1 to localize to forks and its chromatin remodeling activity are required to prevent 53BP1 accumulation on active forks. As shown previously, the ATR-mediated phosphorylation of ATAD5, upon HU induced stalled fork accumulation, interacts with proteins at reversed forks proteins (*39*). We suggest that in the absence of SMARCAD1, enhanced ATAD5-RLC levels causing PCNA dissociation from forks leads to frequent fork stalling and consequently accumulation of reversed forks. 53BP1 binding to stalled /reversed forks further stabilize ATAD5 via their direct interaction which leads to increased PCNA unloading. Upon HU induced fork stalling, the NΔ-SMARCAD1 cells show consistent accumulation of 53BP1-ATAD5 with forks even upon release from HU that further leads to poor PCNA recovery causing delayed fork restart and defective fork progression. The enzymatic activity of SMARCAD1 could be required to displace or reposition 53BP1-bound nucleosomes at regressed arm of reversed forks, similar to previously reported at DSBs (*23*), as the ATPase-dead mutant of SMARCAD1 shows defect in fork progression and restart efficiency similar to NΔ-SMARCAD1. Furthermore, previously it was suggested that the loss of 53BP1 restores HR in SMARCAD1-depleted cells which is responsible for developing resistance to replicative stress-inducing drugs (*23*). However, with this study using separation-of-function SMARCAD1 mutant, which is HR proficient but defective for fork stability, shows that the extent of damage generated upon replication stress is rather responsible for the cellular sensitivity and is not because of unrepaired DSBs due to lack of HR. This further suggests that the role of SMARCAD1 at forks is crucial for tolerance to replication stress inducing agents. We have, therefore, revealed a moonlighting function of SMARCAD1 at the replication forks in displacing 53BP1 to maintain replication fork progression and genome stability. Other NHEJ factors such as RIF1, PTIP etc. have also been found in association with replication forks. Therefore, it would be interesting to investigate if 53BP1 works in complex with NHEJ machinery or have a separate role in association with ATAD5-RLC complex to regulate PCNA homeostasis and thereby fork dynamics.

### An essential role of SMARCAD1 in the viability of BRCA1-defective tumors

BRCA1/2 factors, independent of their role in HR, protect replication forks and prevent their collapse into genome-destabilizing DSBs (*6, 7*). SMARCAD1 has been shown to be epistatic with BRCA1 in the context of HR, (*23, 29*). However, here, we show contrasting differences in role of SMARCAD1 than that of BRCA1 by a) differential enrichment of SMARCAD1 and BRCA1 at the replication forks, where SMARCAD1 preferentially associates with active forks while BRCA1 with stalled forks (Fig. 1A) (8), b) stalled forks induced by 4mM HU in absence of SMARCAD1 are not degraded unlike upon loss of BRCA1, c) loss of SMARCAD1 but not BRCA1 causes defects in unperturbed replication fork progression (Fig. 3A and fig. S5A) (*2*) and finally, d) loss of 53BP1 in BRCA1 deficient cells that restores HR repair capacity, do not rescue sensitivity of BRCA1 mutants to Cisplatin treatment (Fig. 5F) (*60*). However, loss of 53BP1 in SMARCAD1 mutant rescues Cisplatin sensitivity, suggesting replication stress sensitivity is uncoupled from HR repair and that SMARCAD1’s role at active replication forks is distinct from that of BRCA1’s role at stalled replication forks to maintain tolerance towards replication stress inducing agents. Together these data suggest distinct role of SMARCAD1 and BRCA1 at replication forks acting in two independent pathways, where SMARCAD1 mediates active fork stability while BRCA1 mediates stalled fork protection. However, both the pathways are interdependent for maintaining replication fork integrity, which is also conserved across species, from mouse to human (Fig. 6, C and F). Moreover, loss of SMARCAD1 results in enhanced accumulation of DNA damage and ultimately, synthetic lethality in mouse- BRCA1- defective tumors irrespective of their HR status. These findings suggest that these factors may work in parallel to stabilize replication forks and act synergistically to maintain fork integrity. Intriguingly, loss of Mre11 but not 53BP1 rescued fork progression defects that appeared upon loss of both BRCA1 and SMARCAD1 together in cells. This data imply that BRCA1-mediated stabilization of stalled forks allows the enrichment of 53BP1, which further delays fork restart in absence of SMARCAD1. Similarly, loss of SMARCAD1 in BRCA1 deficient mouse tumor organoids could result in Mre11-mediated fork degradation, as observed for human fibroblast cells, which subsequently result in massive accumulation of unrepaired DSBs in genomes, causing synthetic lethality.

In summary, we have shown a conserved interplay between SMARCAD1 and BRCA1 in stabilization of replication forks, where SMARCAD1 stabilizes active forks while BRCA1 protects stalled forks to maintain genome integrity (fig. S6). Notably, SMARCAD1 mediated stabilization of unperturbed forks promotes cellular proliferation in BRCA1-deficient mouse breast tumor, cells and organoids, independently of their HR- and PARPi-resistance status. Similarly, the correlation of reduced chances of survival after chemotherapy in cancer patients with enhanced expression of SMARCAD1 along with reduced expression of BRCA1, suggest that stabilization of active forks promotes tolerance towards chemotherapy in BRCA1-defective tumors. Finally, the observation that SMARCAD1 become essential for genome stability and cellular survival in the absence of BRCA1, suggest that targeting the stability of active replication forks has the potential to be a clinically effective remedy for BRCA-deficient tumors, naïve or chemoresistant. It also suggests that SMARCAD1 could be a strong candidate for development of novel therapeutic treatment for BRCA1-deficient cancer patients.

## AUTHOR CONTRIBUTIONS

C.S.Y.L. conducted all the QIBC, FACS, and PFGE experiments. M.v.T performed iFRAP, chromatin fractionation experiments and with help from Y.Z performed crosslinked chromatin IP experiments. V.G. performed all the fiber experiments and IF experiments related to ATAD5. M.P.D. performed all the cloning experiments and clonogenic assays using mouse tumor cells/organoids under the supervision of J.J. Y.Z. with the help of M.v.d.D. performed cloning experiments of cDNA-SMARCAD1. E.M.M. with help from C.S.Y.L performed clonogenic assays with MRC5 cells and chromatin fractionations for RAD51. H.L. helped C.S.Y.L, and M.v.d.D. in cloning experiments in MRC5 cells. M.v.R, W.Z. and I.S. analyzed fluorescence microscopy data. The assistance to use high-content imaging microscope facility was provided by M.v.R and P.J.F. J.D. analyzed mass-spectrometry data. J.G.S.C.S.G analyzed TCGA ovarian breast cancer data. D.W. analyzed RNA-Seq data. J.A.M supervised the iFRAP and chromatin fractionation experiments. A.R.C supervised the iPOND experiments performed by C.M and EM experiments performed by E.M.M. N.T. conceptualized the project, supervised it, and wrote the manuscript.

## ACKNOWLEDGMENTS

We thank Roland Kanaar, Wim Vermeulen, and Claire Wyman for stimulating discussions and sharing important reagents used in the manuscript; Kyungjae Myung and Kyooyoung Lee for ATAD5 antibody and sharing technical information, Dik van Gent for 53BP1 antibody, Ewa Goggola for help with initial phase of mouse tumor cells culture. This work was supported by grants from the Daniel den Hoed Stiching Young Scientific Talent grant (DDHS#108341) to NT and the Oncode Institute partly financed by the Dutch Cancer Society funded grant (KWF grant 10506) to JAM, Erasmus MC Daniel den Hoed instrument grant to ARC and startup funds from the Erasmus MC to NT.

## DECLARATION OF INTERESTS

The authors declare no competing interests.

### Data and materials availability

**NCBI bioproject accession number:** PRJNA609878

### Supplementary Materials

Materials and Methods

Figures S1-S6

Tables S1-S3

## MATERIALS AND METHODS

### Cell line generation

Plasmid transfections were performed using X-tremeGENE 9 DNA transfection agent (Roche) according to the manufacturer’s protocol. To generate MRC5 NΔ-SMARCAD1 cells, MRC5 WT cells were transfected with pLentiCRISPR-V2 plasmid (addgene: #52961) containing a gRNA sequence targeting exon 2 of SMARCAD1, followed by puromycin selection (1 μg/ml).To generate MRC5 SMARCAD1^-/-^, two gRNA sequences targeting exon 2 and exon 24 of SMARCAD1 were selected and co-transfected with homologues repair template containing mClover reporter gene as fluorescent selection marker for FACS sorting. The primers for gRNA are listed in Table S3. To express mClover-SMARCAD1 full length/ SMARCAD1 K528R mutant cDNA in MRC5 cells, gRNAs targeting SMARCAD1 exon2 and exon 24 were used and co-transfected with mClover-SMARCAD1 full length/ SMARCAD1 K528R mutant cDNA respectively in MRC5 WT and NΔ-SMARCAD1 cells. The K528R mutant was generated using the full length SMARCAD1 cDNA by site-directed mutagenesis. The primer for site-directed mutagenesis are listed in Table S3.

To generate GFP-tagged PCNA knock-in MRC5 cells, a gRNA sequence targeting exon 2 of PCNA was selected and inserted into lentiCRISPR V2 (addgene Plasmid #52961). MRC5 WT and NΔ-SMARCAD1 cells were transfected with the gRNA and the FLAG-GFP-PCNA repair template and sorted by FACS sorting.

### Cell culture

All MRC5 human fibroblasts were cultured in a 1:1 ratio of Dulbecco’s modified Eagle’s medium (DMEM) and Ham’s F10 (Invitrogen) supplemented with 10% fetal calf serum (FCS, Biowest) and 1% Penicillin–Streptomycin (PS, Sigma-Aldrich) at 37 °C and 5% CO_2_ in a humidified incubator.

KB1P-G3, KB1P 177-a5(*46, 47*) and KB1P-G3B1 (*47*) have been described previously. All KB1P mouse tumor cell lines were cultured in DMEM/F12+GlutaMAX (Gibco) containing 5μg/ml Insulin (Sigma-Aldrich), 5 ng/ml cholera toxin (Sigma-Aldrich), 5 ng/ml murine epidermal growth-factor (Sigma-Aldrich), 10% FCS and 1% PS (Sigma-Aldrich) and under low oxygen conditions (3% O_2_, 5% CO_2_ at 37°C).

All tumor-derived organoid lines have been described before(*48*). KB1P4N.1 and KB1P4R.1 tumor organoids were derived from a mammary KB1P PARPi-naïve and PARPi-resistant tumor, respectively (female donor). Cultures were embedded in Culturex Reduced Growth Factor Basement Membrane Extract Type 2 (BME, Trevigen; 40 ml BME:growth media 1:1 drop in a single well of 24-well plate) and grown in Advanced DMEM/F12 (Gibco) supplemented with 1M HEPES (Gibco), GlutaMAX (Gibco), 50 units/ml penicillin-streptomycin (Sigma-Aldrich), B27 (Gibco), 125 mM N-acetyl-L-cysteine (Sigma-Aldrich) and 50 ng/ml murine epidermal growth factor (Sigma-Aldrich). Organoids were cultured under standard conditions (37°C, 5% CO_2_) and regularly tested for mycoplasma contamination. Mouse embryonic stem cells (mESCs) were maintained in 2i media deficient in lysine, arginine, and l-glutamine (PAA) at 37 °C and 5% CO_2_ in a humidified incubator. For SILAC labeling, cells were grown in medium containing 73 µg/ml light [^12^C_6_]-lysine and 42 µg/ml [^12^C_6_, ^14^N4]-arginine (Sigma-Aldrich) or similar concentrations of heavy [^13^C_6_]-lysine and [^13^C_6, N4_ ^15^]-arginine (Cambridge Isotope Laboratories).

## Method Details

### siRNA transfection, shRNA transduction and Cell Titre assay

siRNA transfection was done with lipofectamine RNAiMAX (Thermofisher) according to the manufacturer’s protocol for 2 consecutive days. Knockdown efficiency was checked by immunoblot. Details of siRNA oligomers and shRNAs used in this study are given in Table S3.

Transductions were done in duplicate in KB1P mouse tumor cells. After 3 days of selection, KB1P mouse tumor cells were expanded to 10cm dishes. 5 days post passage, 10cm dishes were fixed with 4% formaldehyde, stained with 0.1% crystal violet, and quantification was carried out by determining the absorbance of crystal violet at 590 nm after extraction with 10% acetic acid.

3D Tumor-derived organoids were transduced according to a previously established protocol(*48*). Puromycin selection was carried out for 3 consecutive days after transduction at a concentration of 3 μg/ml. Pictures were taken at day 5. For quantification, cells were incubated with Cell-Titer Blue (Promega) reagent at day 5.

### Chromatin fractionation

Cells were lysed in lysis buffer (30 mM HEPES pH 7.6, 1 mM MgCl2, 130 mM NaCl, 0.5% Triton X-100, 0.5 mM DTT and EDTA-free protease inhibitor cocktail (Roche)), at 4 °C for 30 minutes. Chromatin-containing pellet was spinned down by centrifugation at 16,000g for 10 minutes and resuspended in lysis buffer supplemented with 250 U/µL of Benzonase (Merck Millipore) and incubated for 15 minutes at 4 °C.

### Live cell confocal imaging

Live cell confocal laser-scanning microscopy was carried out as described before (*61*), with minor adjustments. All live cell imaging experiments were performed using a Leica TCS SP5 microscope (with LAS AF software, Leica) equipped with HCX PL APO CS 63x oil immersion objective (Leica Microsystems), at 37 °C and 5% CO_2_. For Inverse FRAP (iFRAP), GFP-PCNA expressing WT and NΔ-SMARCAD1 MRC5 cells were seeded on 24 mm coverslips. Cells were continuously bleached at high 488 nm laser outside the selected GFP-PCNA foci and the fluorescence decrease of the selected foci was determined over time. The resulting dissociation curves were background-corrected and normalized to pre-bleach values, set at 1.

### DR-GFP reporter assay

The procedure for DR-GFP reporter was described previously (*30*) and applied with minor alterations. After being seeded in a 6-well plate for 24 hours, cells were co-transfected with 1.5 μg of DR-GFP reporter plasmid (addgene #26475) and 1.5 μg I-Scel expression vector (addgene # 26477) or empty vector using X-tremeGENE 9 DNA transfection agent (Roche) according to the manufacturer’s protocol for 24 hours. p-MAX-GFP plasmid (addgene #16007) was transfected in parallel to assess transfection efficiency. Another round of transfection was done on day 2. On day 3, cells were harvested and GFP expression was analyzed by flow cytometer.

### DNA fiber analysis

DNA fiber analysis was carried out according to the standard protocol as mentioned previously (*34*). Briefly, cells were sequentially pulse-labeled with 30 μM CldU (MP Biomedicals) and 250 μM IdU (Sigma-Aldrich) according to the schematic in each figure. For mirin treatment, 100μM Mirin was added to the cell culture media for 2 hours prior to the CldU and IdU labeling. After labeling, cells were collected and resuspended in PBS at 5 × 10^5^ cells per ml. The labeled cells were mixed with equal amount of unlabeled cells, and 2.5 µl of mixed cells were added to 8 µl of lysis buffer (200 mM Tris-HCl, pH 7.5, 50 mM EDTA, and 0.5% (w/v) SDS) on a glass slide. After 8 minutes, the slides were tilted at 30–45°, and the resulting DNA spreads were air dried, fixed in 3:1 methanol/acetic acid overnight at 4 °C. The fibers were denatured with 2.5 M HCl for 1 hour, washed with PBS and blocked with 0.1% Tween 20 in 2% BSA/PBS for 40 minutes. The newly replicated CldU and IdU tracks were labeled for 2 hours in dark with anti-BrdU antibodies recognizing CldU (1:100)(Abcam, ab6326) and IdU (1:100)(BD, 347580), followed by 1 hour incubation with secondary antibodies in the dark: anti–mouse Alexa Fluor 488 (1:250) (Invitrogen, A-11001) and anti–rat Cy3 (1:250) (Jackson Immuno-Research Laboratories, 712-166-153). Fibers were visualized and imaged by Carl Zeiss Axio Imager D2 microscope using 63X Plan Apo 1.4 NA oil immersion objective. ImageJ software was used for the quantification. The Kruskal-Wallis test followed by Dunn’s multiple comparison test was applied for statistical analysis using the GraphPad Prism Software. The combined summary of DNA fiber spread data analysis is given in Table S2.

### Immunoblot and antibodies

After lysed with RIPA buffer supplemented with protease inhibitor (Roche)(whole cell lysate) or resuspended in chromtain fractionation lysis buffer (chromatin bound proteins), samples were mixed with 2x Laemmli sample buffer (Supelco) and heated at 95°C for 5 minutes. Samples were loaded on 4–12% NuPAGE Bis-Tris Gel (Novex life technologies) and transferred to a Polyvinylidene difluoride (PVDF) membrane (0.45μm, Immobilon). Membranes were blocked with 5% BSA in PBS for 1 hour at room temperature and incubated with mouse anti-alpha-tubulin monoclonal antibody (sigma, T6074), rabbit anti-SMARCAD1 antibody (Atlas, HPA016737), mouse anti-PCNA monoclonal antibody (abcam, ab29), rabbit anti-Histone H1.2 antibody (abcam, ab17677), mouse anti-RPA32/RPA2 antibody (abcam, ab2175), rabbit anti-GFP antibody (abcam, ab290), rabbit anti-Atad5 antibody (abcam, ab72111), rabbit anti-Histone H3 antibody (abcam, ab1791) or mouse anti-p37 (GeneTex, 1320) diluted in blocking buffer overnight at 4°C. Membranes were washed in 0.1% Tween-20 in PBS on the following day, followed by incubation with secondary antibody coupled to near-IR dyes CF™680/CF™770 (1:10,000)(Sigma, SAB4600205 &SAB4600215). Antibodies were visualized using an Odyssey CLx infrared scanner (LiCor). ImageJ software was used for the quantification of bands on western blots, wherever applicable.

### Immunofluorscence staining

Cells were labeled with EdU (10 µM) for 30 minutes to identify cells in S-phase, unless otherwise mention for the EdU progression experiment. For HU treated samples, EdU is labeled before the HU treatment. For analysis of the chromatin bound protein, cells were first pre-extracted with 0.1% Triton-X 100 in ice-cold CSK buffer for 5 minutes at 4°C before fixation. Cells are fixed in 2% formaldehyde in PBS for 15 minutes at room temperature for SMARCAD1 (Atlas, HPA016737), 53BP1 (Novus, NB100-304), RAD51 (B-Bridge International, 70-001) and γH2AX (Merck Millipore, 05-636) or 100% -20°C methanol for 10 minutes for PCNA(abcam, ab29). Subsequently, samples were permeabilized in 0.1% Triton-X 100 in PBS for 10 minutes, and blocked with 5% BSA in PBS. Samples were subsequently stained with primary antibody diluted in blocking buffer, followed by incubation in fluorescence conjugated secondary antibody. EdU was visualized with a click-it reaction using Alexa Fluor® 488 azide (Invitrogen, C10337) or Alexa Fluor® 594 azide (Invitrogen, C10646) according to the manufacturer’s protocol. Samples were washed with PBS and incubated with 0.1ug/ml DAPI for 15 minutes. ProLong™ Gold antifade mountant (Invitrogen) was used to mount the samples on the glass slides for coverslip samples.

### Image acquisition and image analysis

Coverslip images were obtained using a LSM700 microscope equipped with a plan-apochromat 63x/1.4 Oil M27 objective (Carl Zeiss Micro imaging), or SP5 microscope equipped with HCX PL APO CS 63x Oil objective (Leica). The analysis of the image data has been conducted using custom-built ImageJ plugins. The detection of EdU positive (and negative) cells was performed using the 488 nm channel in combination with the DAPI channel by applying a cross entropy based thresholding and the binary watershed segmentation (in order to deal with touching cells). The adjustment of brightness and contrast was applied differently due to differential backgrounds in the indicated cell lines of Fig. 1G for the qualitative representation. To compute the Pearson and Manders’ overlap coefficients in Fig. S4B, the 53BP1 foci in 488 and 568 nm channels for EdU positive cells were segmented using an à-trous wavelet transform with 3 scales, and the wavelet coefficients were thresholded at the level of 3-sigma (*62*). To measure the distance between 53BP1 and EdU foci in Fig. 5A, a line of 3μm was drawn across the proximal foci and the intensity of the two channels were measured using multi plot in imageJ. Further analysis was done using Microsoft Excel. For high-content imaging given in Fig. 1B, 1C, 2C, 4A, 4C, 5E & fig. S1D and S2G, all the data was obtained using Opera Phenix High-Content Screening System (PerkinElmer) with 40x water objective (NA 1.1) and analyzed with the Harmony v4.9 high-content imaging and analysis software (PerkinElmer) using a custom script. At least 75 field per well were imaged as a Z-stack of 8 planes (stepsize 1μm). In the maximum projection, nuclei were detected using the DAPI signal and filtered for nuclear roundness (>0.7) and size (70 250 μm^2^) to exclude dead nuclei, and clusters of multiple nuclei. Selection of S phase cells was based on EdU signal in UT and HU block condition. In HU release condition, S phase cells were determined by intensity of PCNA median. The pixel intensities (sum) were determined in the DAPI, 488 nm and 568 nm channel for the individual nucleus. PCNA sum normalized to DAPI sum was shown in the bar chart. For quantification of EdU positive foci in Fig. 1B & 1C and fig. S1D, an additional mask was generated based on the detection of local intensity maxima (region to spot intensity) in the EdU channel, and used for quantification of spot intensities together with spot contrast in the 488 & 568 nm channels. For quantification of RAD51 positive foci in Fig. 2C, a mask was generated in the RAD51 channel using the detection of local intensity maxima (region to spot contrast and intensity) in the RAD51 channel, with an upper threshold for spot radius. The desired quantified values for each foci/cell were exported to the Tibco spotfire software for generation of scatter diagrams.

### RNA extraction, Reverse Transcription, Real-time qPCR and RNA-seq

Total RNA was extracted using the ReliaPrep™ RNA Miniprep Systems (Promega) according to the manufacturer’s instructions. 1000 ng of total RNA was used to synthesis cDNA using M-MLV Reverse Transcriptase, RNase H Minus, Point Mutant (Promega) according to the manufacturer’s instructions. Real-time qPCR was performed using the GoTaq^®^qPCR Master Mix (Promega), beta-actin was used for normalization. Primers used for qPCR are listed in Table S3.

NGS short reads were trimmed using fastp and processed using Kalliso, an RNAseq quantification program that uses a pseudoalignment method of assigning reads to genomic locations in lieu of a more costly traditional alignment(*63*). The human transcriptome, version GRCh38.p12, was indexed, the paired, trimmed reads assigned to transcripts, and read counts converted to transcripts per million (TPM) by Kallisto. TPMs from transcripts originating from the same gene were aggregated and relative expression levels were computed as the log2 fold change relative to the matched wild type using an in-house script (available upon request). RPKM values were computed from TPMs using the median transcript length per gene. Pseudoalignments, output by Kallisto in standard BAM format, were used to assess transcript structure such as the assignment of the transcription start for NΔ-SMARCAD1. Boxplots and barplots were produced using ggpubr and ggplot2 respectively in R program (the R Foundation).

### iPOND-SILAC mass-spectrometry

For iPOND experiments, light lysine and arginine labeled mESCs cells were incubated with 10 µM EdU for 10 minutes and treated with 4mM HU (Sigma-Aldrich) for 3 hour to stall the DNA replication forks. Heavy lysine and arginine labeled mESCs cells were only incubated with 10 µM EdU for 10 minutes. After labeling and treatment cells were cross-linked with 1% formaldehyde for 10 min at room temperature, quenched with 0.125 M glycine, washed with PBS and harvested using cell scrapper. Samples were then treated with click reaction containing 25 µM biotin-azide, 10 mM (+) sodium l-ascorbate and 2 mM CuSO_4_ and rotated at 4 °C for 1 h. Samples were then centrifuged to pellet down the cells; supernatant was removed and replaced with 1 ml Buffer-1 (B1) containing 25 mM NaCl, 2 mM EDTA, 50 mM Tris–HCl, pH 8.0, 1% IGEPAL and protease inhibitor and rotated again at 4 °C for 30 min This step was repeated twice. Samples were centrifuged to pellet down the cells; supernatant was removed and replaced with 500 μl of B1 and sonicated using a Bioruptor Sonicator (Diagenode) using cycles of 20 s ON, 90 s OFF for 30 times at high amplitude. Samples were centrifuged, and supernatant was transferred to fresh tubes and incubated for 1 hour with 200 μl of Dynabeads MyOne C1 (Sigma-Aldrich) for the streptavidin biotin capture step. Proteins were eluted, and mass-spectrometry was performed. At least two peptides were required for protein identification. Quantitation is reported as the log_2_ of the normalized heavy/light ratios. SILAC data were analyzed using MaxQuant. The resulting output tables of two independent experiment were merged and used as the input for calculating the average fold-change to identify significantly upregulated proteins in unperturbed forks and stalled forks based on H:L ratio in the SILAC experiment in the MaxQuant software (*9*).

### Crosslinked immunoprecipitation

The procedure for in vivo crosslink and immunoprecipitation was described previously(*61*) and applied with minor alterations. After removal of medium, cells were cross-linked in 1% formaldehyde in serum-free medium for 10 minutes at room temperature. Crosslinking reaction was stopped with 0.125 M of glycine and cells were collected in ice cold PBS supplemented with 10% glycerol. Crosslinked cells were scrapped and chromatin was purified as described(*61*). Chromatin was sheared using a Bioruptor Sonicator (Diagenode) using cycles of 20 s ON, 60 s OFF during 15 minutes, after which samples were centrifuged. The supernatant containing crosslinked chromatin was used for immunoprecipitation. For immunoprecipitation, extracts were incubated with either GFP-trap beads (ChromoTek), 53BP1 (1.8μg) or SMARCAD1 (1.8μg) antibody overnight at 4 °C. For IP with 53BP1 and SMARCAD1 antibody, Protein A agarose/Salmon Sperm DNA slurry (Millipore) was added for 4 hour at 4 °C. Subsequently, beads were washed five times in RIPA buffer and elution of the precipitated proteins was performed by extended boiling in 2x Laemmli sample buffer (Sigma-Aldrich) for immunoblotting analysis.

### Clonogenic survival assay

Cells were seeded in triplicate in 10cm culturing dish and treated with a single dose of olaparib (selleckchem), cisplatin (Sigma-Aldrich) or hydroxyurea(HU) (Sigma-Aldrich) 1 day after seeding. For hydroxyurea, HU was given at the indicated concentration for 24 hours or 48 hours as indicated in the Fig. legend before being washed off and replaced with new medium. For olaparib, different concentrations of olaparib were given to the cells throughout the whole experimental process. For cisplatin, different concentrations of cisplatin were given to the cells for 4 hours before being washed off and replaced with new medium, except the 1 μM cisplatin group in Fig. 5F and S3H, which were given throughout the whole experimental process. After 1 week, colonies were fixed and stained in a mixture of 43% water, 50% methanol, 7% acetic acid and 0.1% Brillant Blue R (Sigma-Aldrich) and subsequently counted with the Gelcount (Oxford Optronix). The survival was plotted as the mean percentage of colonies detected following the treatment normalised to the mean number of colonies from the untreated samples.

### Cell cycle analysis

Cells were grown to 70–80% confluency in a 10cm culturing dish. Cells were labeled with EdU for 30 minutes followed by fixation for 10 minutes in 4% formaldehyde in PBS at room temperature. Cells were then washed with 1% BSA/PBS and permeabilized in 0.5% saponin buffer in 1% BSA/PBS. Incorporated EdU were labelled with the click-it reaction using Alexa Fluor® 594 azide according to the manufacturer’s protocol (Invitrogen). DAPI was used to stain the DNA.

### Electron microscope analysis

EM analysis was performed according to the standard protocol(*35*). For DNA extraction, cells were lysed in lysis buffer and digested at 50 °C in the presence of Proteinase-K for 2 hour. The DNA was purified using chloroform/isoamyl alcohol and precipitated in isopropanol and given 70% ethanol wash and resuspended in elution buffer. Isolated genomic DNA was digested with PvuII HF restriction enzyme for 4 to 5 hour. After the digestion, the DNA solution was transferred to a Microcon DNA fast flow centrifugal filter. The filter was washed with TE buffer after spinning for 7 minutes. The benzyldimethylalkylammonium chloride (BAC) method was used to spread the DNA on the water surface and then loaded on carbon-coated nickel grids and finally DNA was coated with platinum using high-vacuum evaporator MED 010 (Bal Tec). Microscopy was performed with a transmission electron microscope FEI Talos, with 4 K by 4 K CMOS camera. For each experimental condition, at least 200 replication fork intermediates were analyzed from three independent experiments and MAPS software (Thermo Fisher) was used to analyze the images.

### Pulsed-field gel electrophoresis

For HU treated samples, cells were treated with 4mM HU for 3 hours, follow or not with a 16 hour release, before harvest for PFGE assay. DSB detection by PFGE was done as reported previously (*9*). Briefly, cells were cast into 0.8% agarose plug (2.5 x 10^5^ cells/plug), digested in lysis buffer (100 mM EDTA, 1% sodium lauryl sarcosine, 0.2% sodium deoxycholate, 1 mg/ml proteinase-K) at 37 °C for 48 hour, and washed in 10 mM Tris-HCl (pH 8.0)–100 mM EDTA. Electrophoresis was performed at 14°C in 0.9% pulse field-certified agarose (Bio-Rad) using Tris-borate-EDTA buffer in a Bio-Rad Chef DR III apparatus (9 h, 120°, 5.5 V/cm, and 30- to 18-s switch time; 6 h, 117°, 4.5 V/cm, and 18- to 9-s switch time; and 6 h, 112°, 4 V/cm, and 9- to 5-s switch time). The gel was stained with ethidium bromide and imaged on Uvidoc-HD2 Imager. ImageJ software was used for the quantification of broken DNA normalized to unbroken DNA for each lane.

### Purification of SMARCAD1 and mass spectrometry

NΔ-SMARCAD1 protein was purified from whole cell lysate using MRC5 NΔ-SMARCAD1 cell line. Cells were resuspended in the IP buffer and sheared 10 time as 15s on and then 45s off at mode High using a Bioruptor Sonicator (Diagenode) at 4°C, and incubated with 500U of Benzonase (Merck Millipore) for 60 minutes, after which samples were centrifuged. The supernatant was used for immunoprecipitation. For immunoprecipitation, extracts were incubated with SMARCAD1 (1.8μg) antibody overnight at 4 °C. Protein A agarose/Salmon Sperm DNA slurry (Millipore) was added for 2 hour at 4 °C. Subsequently, beads were washed five times in IP buffer and elution of the protein was performed by extensive boiling in 2x Laemmli sample buffer (Sigma-Aldrich). Eluted protein was run on 4–12% NuPAGE Bis-Tris Gel (Novex life technologies), gel slices were trypsinized, and peptides were analyzed by mass spectrometry to determine the protein sequence as described previously(*61*).

### Bioinformatic analysis on TCGA datasets

Disease-free survival curves of TCGA high grade serous ovarian carcinoma (HGSOC) patients were generated by the Kaplan–Meier method and differences between survival curves were assessed for statistical significance with the log-rank test. We divided the TCGA ovarian carcinoma patients expressing replication stress markers (CCNE1 overexpression, CDKN2A low expression and/or RB1 deletion) into cohorts according to their BRCA1 mRNA expression levels: BRCA1 low (below median), and BRCA1-high (above median) (*64*). In each of these cohorts, we analysed the correlation between SMARCAD1 expression with outcome. Normalization of expression values was performed using z-score transformation, such that low SMARCAD1 expression with z-score < 0.75 and high SMARCAD1 expression with z-score > 0.75 (fig. S5C). Cohort with BRCA1-high, SMARCAD1-low expression, n = 66; BRCA1-low, SMARCAD1-high expression, n = 10. Cohort with BRCA1-low, SMARCAD1-low expression, n = 87; BRCA1-low, SMARCAD1-high expression n = 10.

### Quantification and Statistical Analysis

For all data, the means, S.D. and S.E.M. were calculated using either Microsoft Excel or GraphPad Prism 8.

---

**Figure S1.**
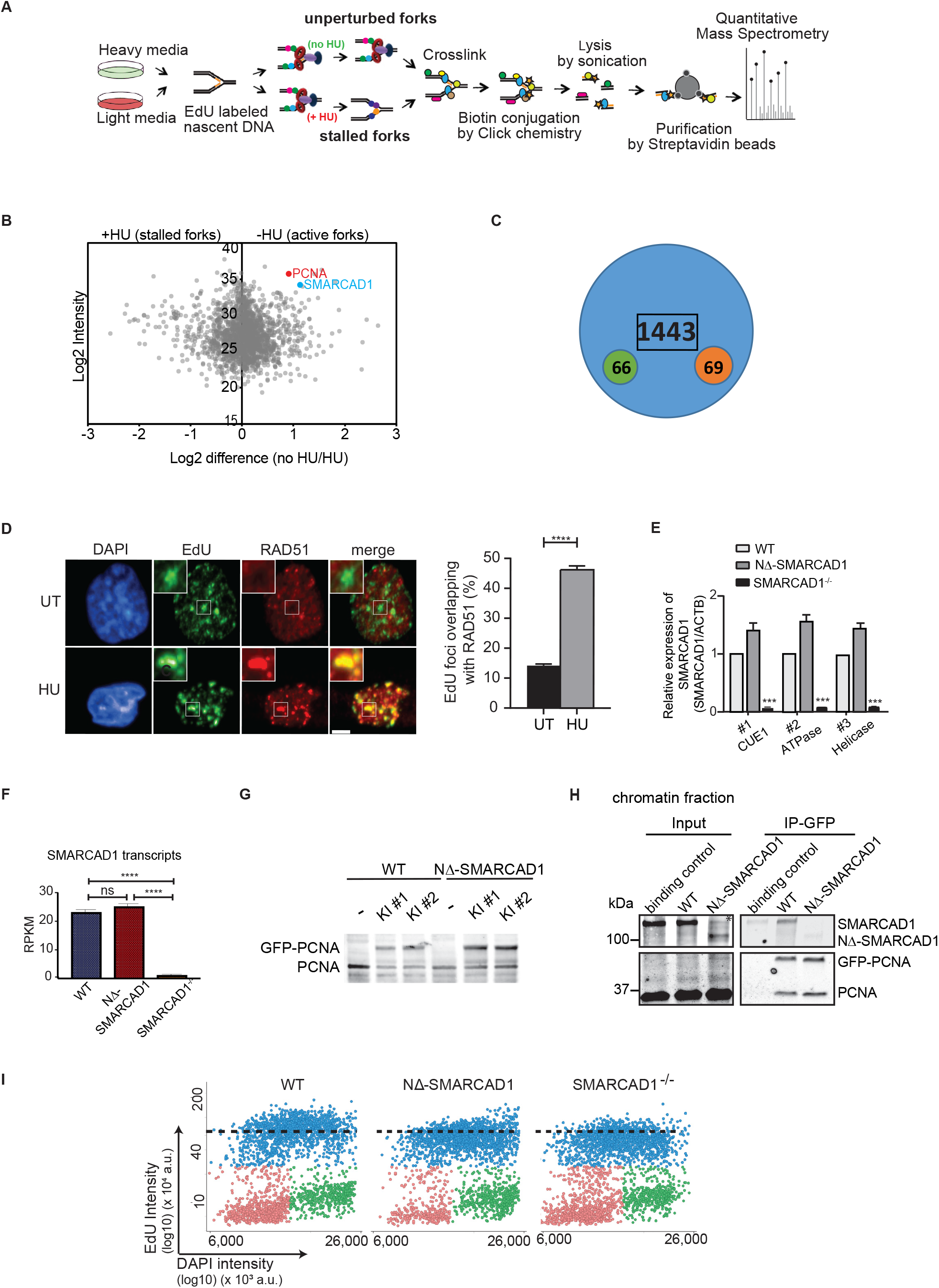
iPOND reveals that SMARACAD1 unlike RAD51 is enriched at unperturbed forks. **(A)**Schematic representation of the iPOND-SILAC-MS experiment. **(B)**Volcano plot showing the distribution of iPOND-SILAC-MS results for average fold-change to identify significantly upregulated proteins in unperturbed conditions based on H:L (no HU: HU) ratio in the SILAC experiment. SMARCAD1 (indicated in blue) and PCNA (indicated in red) show higher enrichment in unperturbed condition. **(C)**Total number of proteins **identified** from two independent iPOND-SILAC-MS experiments using mouse ESCs. Green and red circles represent number of proteins upregulated in unperturbed conditions and HU stalled replication forks respectively. **(D)**(Left) Representative images showing the co-localization of RAD51 (red) to sites of DNA replication as marked by EdU (green) in the presence or absence of HU in human fibroblast MRC5 cells using high-content microscopy (scale bar = 5μm). (Right) Bar chart representing the percentage of EdU foci colocalizing with RAD51 in untreated and 3 hour 4mM HU block condition. (****P ≤ 0.0001, unpaired t-test). **(E)**Transcript levels of SMARCAD1 relative to ACTB in WT, NΔ-SMARCAD1 and SMARCAD1^-/-^ are determined by qRT-PCR and shown as the mean + S.D. (n=3). The normalized value of expression in WT for each primer pair #1, #2 and #3 designed for the exons spanning CUE1, ATPase and Helicase domain, respectively is set to 1. **(F)**Quantification of SMARCAD1 transcript using transcriptome analysis in WT, NΔ-SMARCAD1 and SMARCAD1^-/-^ cells. (n=2) **(G)**Immunoblot showing the GFP-PCNA and PCNA in heterozygous GFP-tagged PCNA knock-in (KI) MRC5 WT and NΔ-SMARCAD1 cells. **(H)**Crosslinked immunoprecipitation of GFP-tagged PCNA expressing endogenously in WT and NΔ-SMARCAD1 cells, using GFP antibody. Western blot analysis was performed using antibodies against PCNA and SMARCAD1. The failure to detect GFP-PCNA band by mouse monoclonal (PC10) antibody mainly in inputs of crosslinked-IP conditions is possibly due to epitope masking under distinct buffer compositions in contrast to IP conditions. The GFP-PCNA band can be easily detected using this antibody in the whole cell extracts prepared in RIPA buffer, as shown in Fig. S1**G**. **(I)**Quantitative image-based cytometry single-cell analysis (QIBC) of EdU labeled WT, NΔ-SMARCAD1 and SMARCAD1-/- cells. G0-1, S and G2/M phase cells are labeled in red, blue and green respectively. Dotted lines represent the mean EdU intensity in WT S-phase cells.

**Figure S2.**
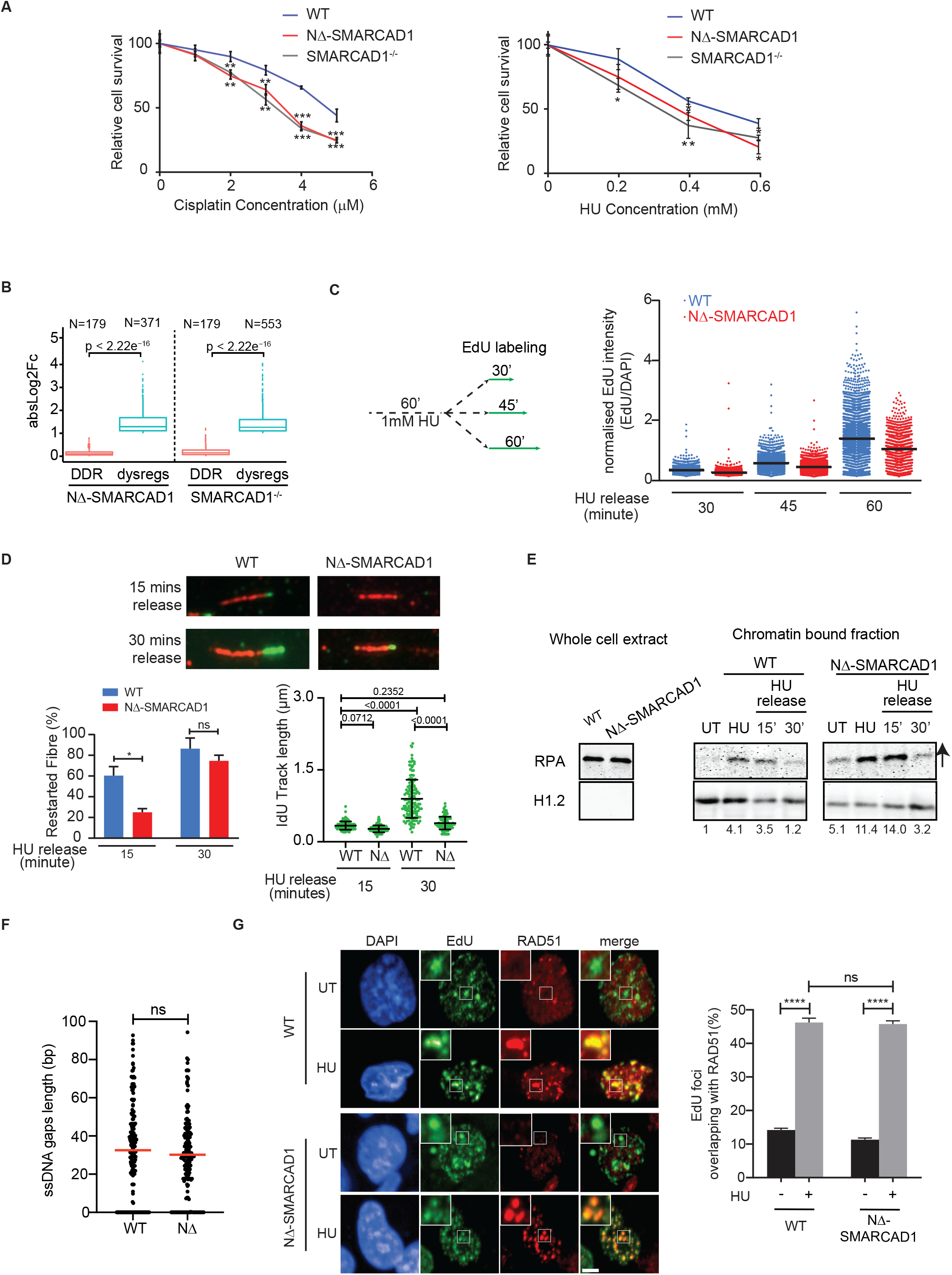
SMARCAD1 is required for efficient fork restart and genome stability. **(A)**Quantification of colony survival assay in WT, NΔ-SMARCAD1 and SMARCAD1^-/-^ cells treated with the indicated concentrations of (left) cisplatin and (right) hydroxyurea. (***P ≤ 0.001, **P ≤ 0.01, *P ≤ 0.05, unpaired t-test) **(B)**Fold change in transcript levels of DNA damage repair (DDR) genes (red) and dysregulated genes (blue) in NΔ-SMARCAD1 and SMARCAD1^-/-^ normalized to WT. **(C)**(Left)Schematic showing the HU release condition with EdU labeling in WT and NΔ-SMARCAD1 cells. (Right) Quantification of EdU intensity by QIBC in >1000 S-phase cells in the HU release conditions for WT and NΔ-SMARCAD1 cells. Cells treated with 1mM HU for an hour were released in EdU containing media for the indicated time before fixation. **(D)**Top panel: representative images showing DNA fibre with IdU track after HU release in WT and NΔ-SMARCAD1 cells. (scale bar = 1μm). Bottom panel: (Left) Bar plot of the percentage of restarted fibres after HU release for 15 and 30 minutes. (*P ≤ 0.05, unpaired t-test). (Right) IdU track length of restarted fibres after HU release for 15 and 30 minutes. (unpaired t-test). **(E)**Immunoblot showing the whole cell extract and chromatin bound fraction of RPA in untreated, HU block and HU release conditions in WT and NΔ-SMARCAD1 cells. Numbers below indicate the quantification of RPA band after normalisation to the loading control. Arrows indicate the position of omitted well between the lanes. **(F)**Quantification of the length of ssDNA gaps at the fork measured by EM. (n=3) (ns, non-significant, unpaired t-test). **(G)**(Left) Representative images showing the co-localization of RAD51 (red) to sites of DNA replicationas marked by EdU (green) in the presence or absence of HU in human fibroblast MRC5 WT and NΔ-SMARCAD1 cells using high-content microscopy. (scale bar = 5μm). (Right) Quantification of percentage of EdU foci overlapping with RAD51 foci.(****P ≤ 0.0001, ns, non-significant, unpaired t-test)

**Figure S3.**
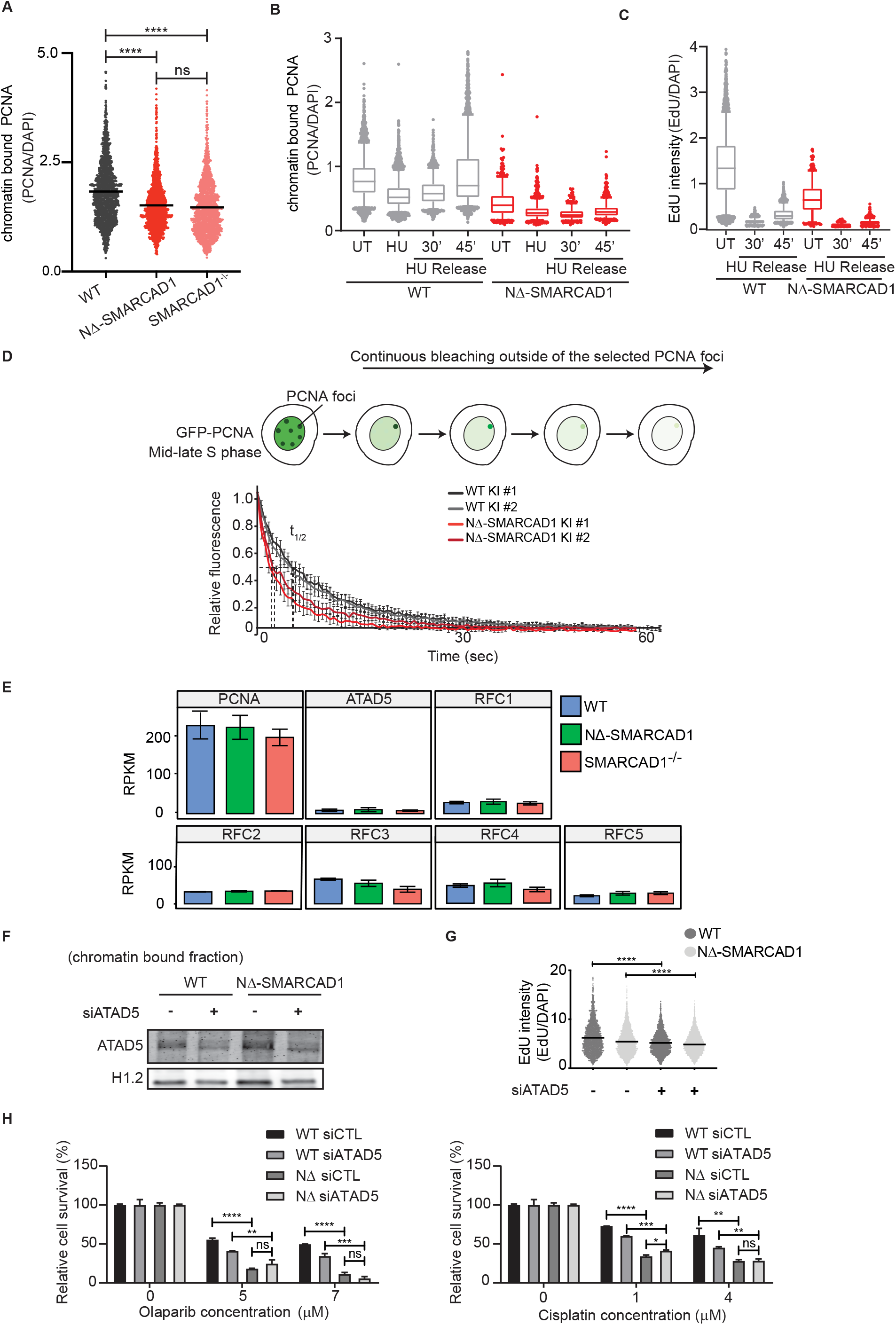
SMARCAD1 maintains PCNA level at replication forks. (**A**) Dotplot of chromatin bound PCNA intensity (normalised to DAPI) in WT, NΔ-SMARCAD1 and SMARCAD1^-/-^ cells. Mean PCNA intensity is indicated. (**B-C**) Boxplot representation of (**B**), chromatin bound PCNA intensity (normalised to DAPI) and (**C**) EdU (normalised to DAPI) upon HU treatment in WT, NΔ-SMARCAD1 cells, corresponding to QIBC analysis shown in Fig. 4C, >1000 S-phase cells were plotted in each condition. **(D)**Top: Schematic for inverse fluorescence recovery after photobleaching (iFRAP) experiment in GFP-tagged PCNA knock-in (KI) cells. Bottom: sample curves for GFP-PCNA in WT and NΔ-SMARCAD1 cells from one experiment (n>12 cells for each experiment, with two independent experiments, mean±2xS.E.M.) **(E)**Quantification of PCNA, ATAD5 and RFC1-5 transcript using transcriptome analysis in WT, NΔ-SMARCAD1 and SMARCAD1^-/-^ cells. **(F)**Immunoblot showing the chromatin bound ATAD5 level in WT and NΔ-SMARCAD1 cells treated with control or ATAD5 siRNA. **(G)**Dotplot of EdU (normalised to DAPI) in WT and NΔ-SMARCAD1 cells treated with control or ATAD5 siRNA. (****P ≤ 0.0001, unpaired t-test). **(H)**Quantification of colony survival assay in WT and NΔ-SMARCAD1 cells treated with control or ATAD5 siRNA and with the indicated concentrations of (left) olaparib and (right) cisplatin. (****P ≤ 0.0001, ***P ≤ 0.001, **P ≤ 0.01, *P ≤ 0.05, ns, non-significant, unpaired t-test).

**Figure S4.**
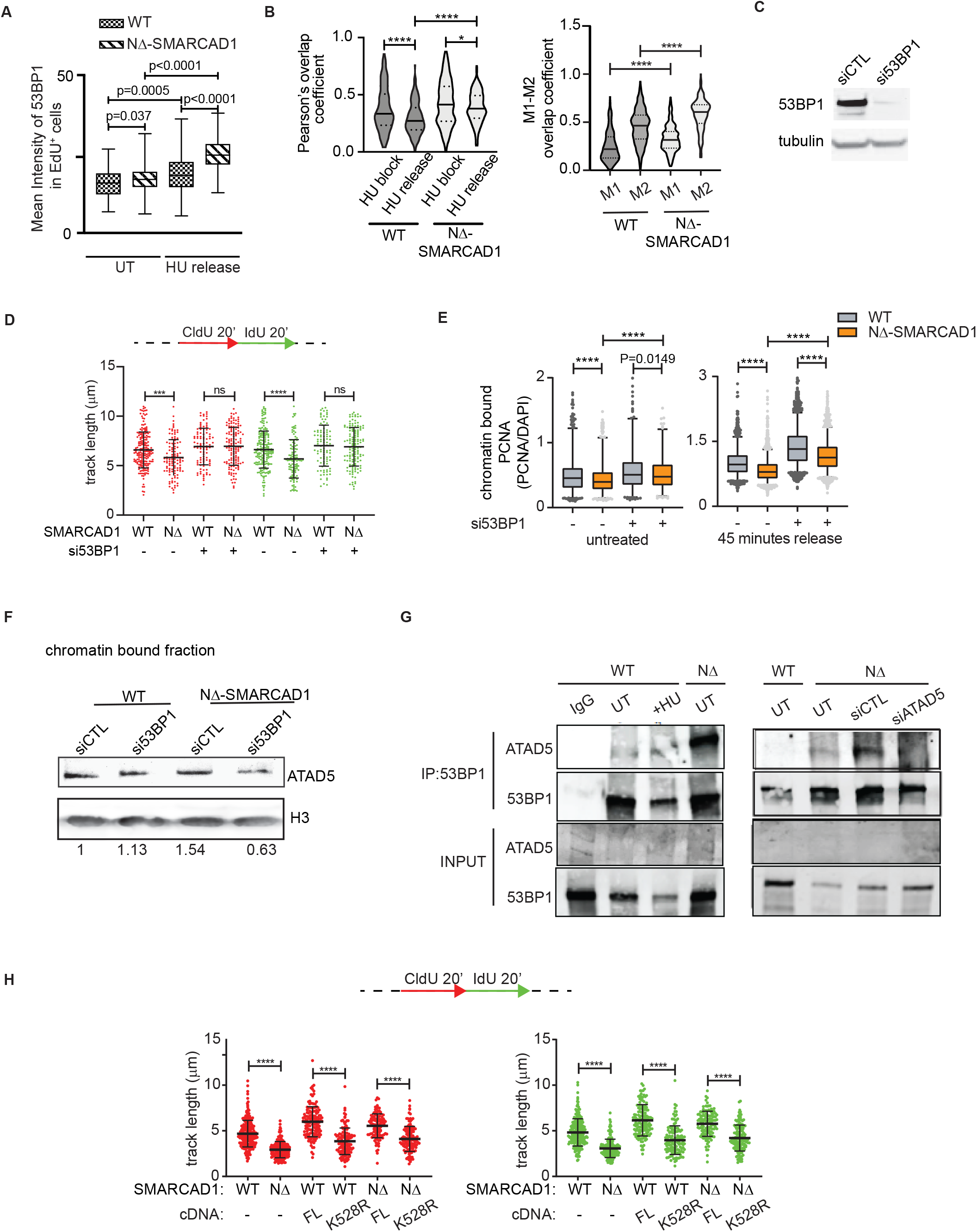
SMARCAD1 precludes 53BP1 enrichment at forks to maintain PCNA levels and facilitate fork progression. **(A)**Boxplot showing mean intensity of 53BP1 in EdU positive WT and NΔ-SMARCAD1 cells in untreated condition and 60 minutes after release from HU treatment (4mM HU for 3 hour). **(B)**Left: Pearson’s overlap coefficient between 53BP1 and EdU in WT and NΔ-SMARCAD1 cells in HU block condition and 60 minutes release after HU treatment (4mM HU for 3 hour). Right: Manders’ M1-M2 overlap coefficients between 53BP1 and EdU in WT and NΔ-SMARCAD1 cells after 60 minutes release from HU treatment (4mM HU for 3 hour). (****P ≤ 0.0001, *P ≤ 0.05, unpaired t-test) **(C)**Immunoblot showing the 53BP1 level in WT cells treated with control or 53BP1 siRNA. **(D)**Top: Schematic for replication fork progression assay with CldU and IdU labeling. Bottom: CldU (red) and IdU (green) track length (μm) distribution for the indicated conditions.(***P ≤ 0.001,****P ≤ 0.0001, ns, non-significant, Kruskal-wallis followed with Dunn’s multiple comparison test, n= 3 independent experiments with similar outcomes.) **(E)**Boxplot showing the intensity of chromatin bound PCNA in EdU positive cells of WT and NΔ-SMARCAD1 in (left) untreated condition and (right) 45 minutes after release from HU treatment (1mM for 1hour), corresponding to QIBC analysis shown in Fig. 5E. (****P ≤ 0.0001, unpaired t-test) **(F)**Immunoblot showing the chromatin bound fraction of ATAD5 levels in WT and NΔ-SMARCAD1 cells upon si-control and si-53BP1 conditions. H3 is used as a loading control. The numbers below the blots show the fold change of ATAD5 after normalisation with H3 relative to WT. **(G)**Crosslinked immunoprecipitation of WT and NΔ-SMARCAD1 cells with the indicated conditions, using 53BP1 antibody. Western blot analysis was performed using antibodies against ATAD5 and 53BP1. **(H)**Top Panel: Schematic for replication fork progression assay with CldU and IdU labeling. Bottom panel: CldU (red) and IdU (green) track length (μm) distribution in cells with/without full length (FL) and K528R ATPase dead cDNA-SMARCAD1 knock-in in WT and NΔ-SMARCAD1 cells. (****P ≤ 0.0001 Kruskal-wallis followed with Dunn’s multiple comparison test, n= 3 independent experiments with similar outcomes)

**Figure S5.**
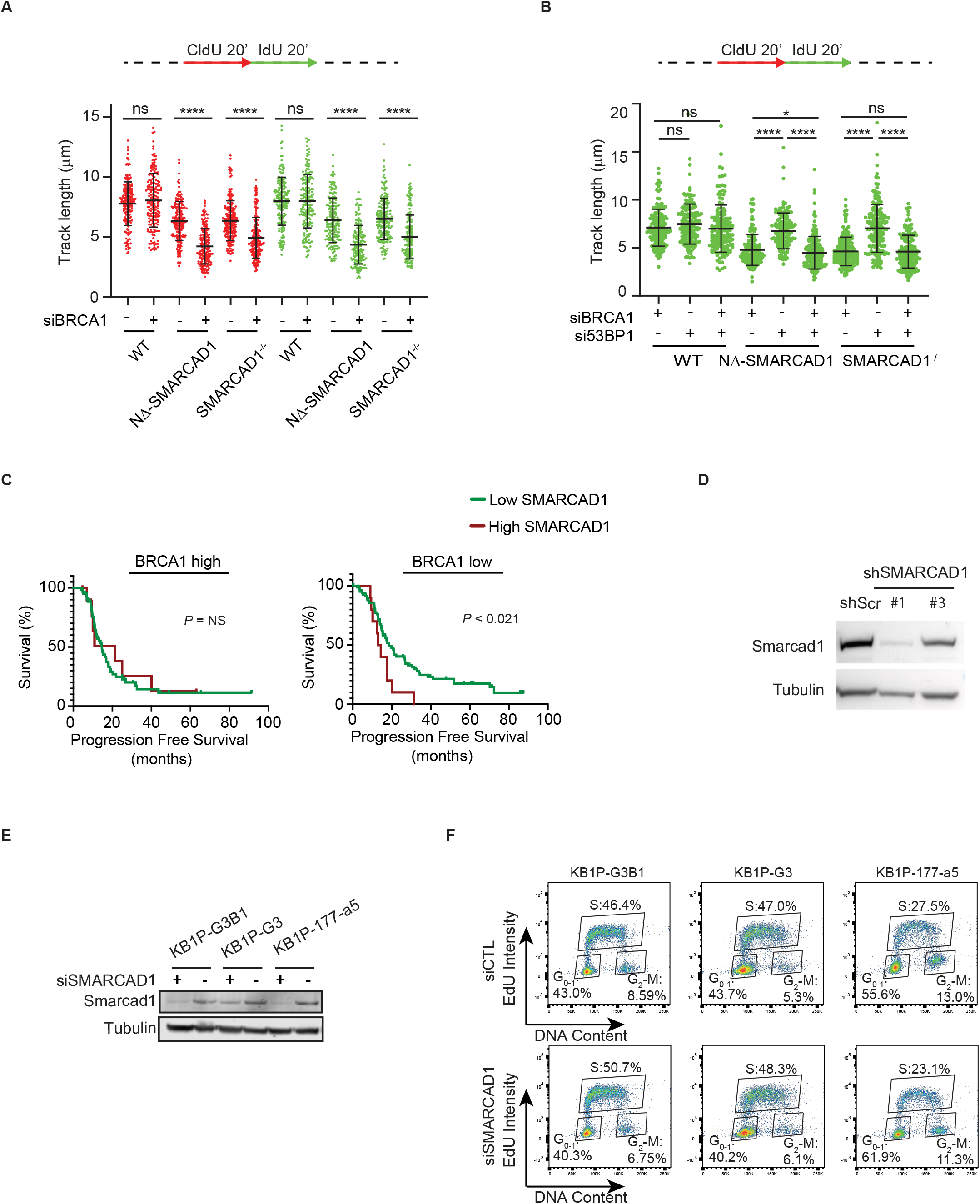
Smarcad1 is essential for proliferation of BRCA1 deficient mouse tumor cells. **(A)**Top: Schematic for replication fork progression assay with CldU and IdU labeling. Bottom: Fork progression assay showing the CldU (red) and IdU (green) track length (μm) distribution for WT, NΔ-SMARCAD1 and SMARCAD1^-/-^ cells treated with si-control or si-BRCA1. (****P ≤ 0.0001, ns, non-significant, Kruskal-wallis followed with Dunn’s multiple comparison test, n= 3 independent experiments with similar outcomes). **(B)**Top: Schematic for replication fork progression assay with CldU and IdU labeling. Bottom: Fork progression assay showing the IdU (green) track length (μm) distribution for WT, NΔ-SMARCAD1 and SMARCAD1^-/-^ cells treated with si-control, si-BRCA1, si-53BP1 or both si-BRCA1 and si-53BP1. (****P ≤ 0.0001, *P ≤ 0.05, ns, non-significant, Kruskal-wallis followed with Dunn’s multiple comparison test, n= 3 independent experiments with similar outcomes). **(C)**Progression-free survival after platinum chemotherapy of ovarian carcinoma TCGA patients with either BRCA1-high or BRCA1-low expression. **(D)**Immunoblot showing the Smarcad1 level in KB1P (Brca1^-/-^; P53^-/-^) tumor cells treated with control (scramble) or two shRNAs (#1 and #3) against Smarcad1. **(E)**Immunoblot showing the Smarcad1 level in KB1P (Brca1^-/-^; P53^-/-^) tumor cells after two days of transfection with FLUC (si-control) or si-SMARCAD1. **(F)**Cell cycle profile of KB1P (Brca1^-/-^; P53^-/-^) tumor cells shown in (**E**).

**Figure S6.**
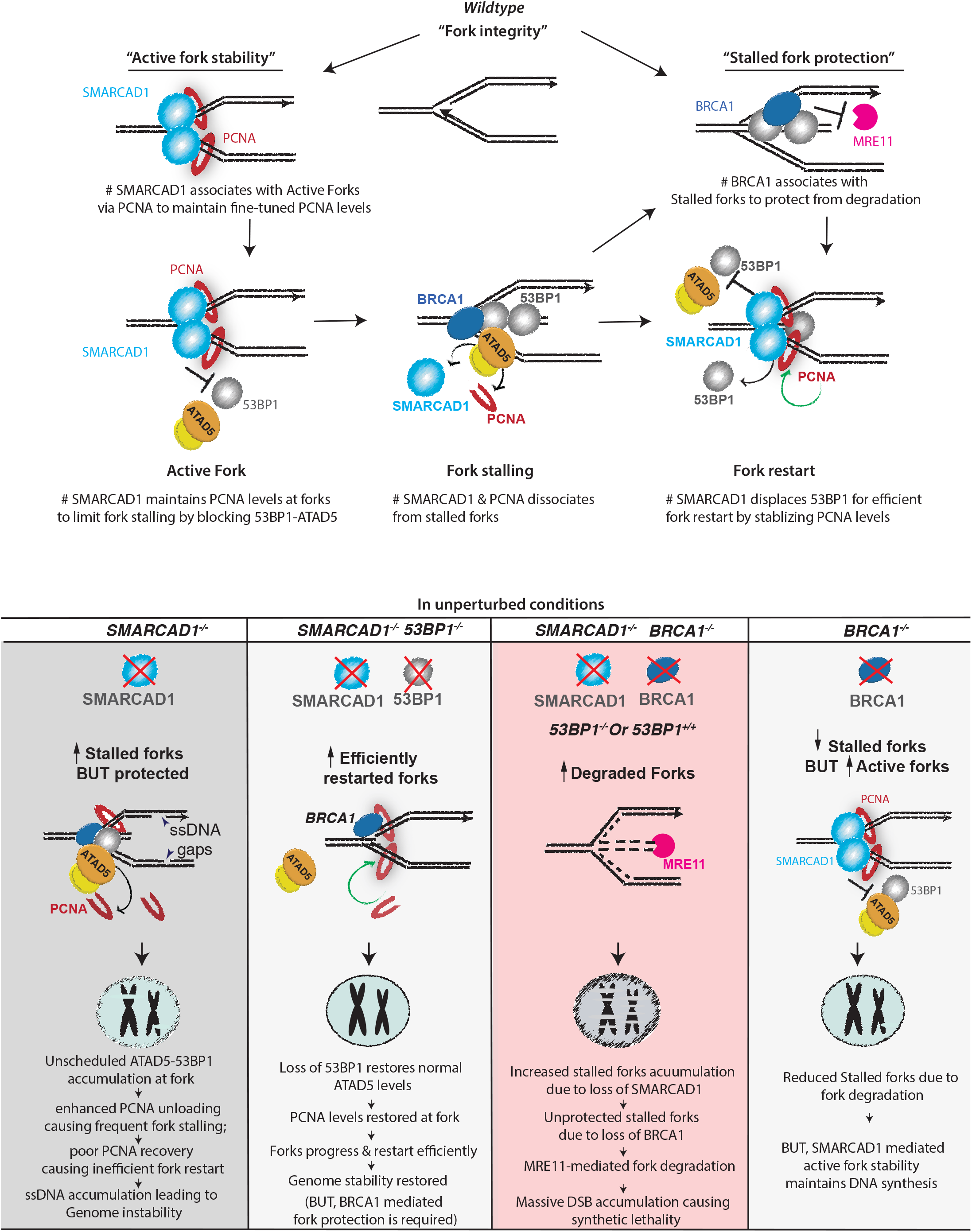
Schematic model depicting the mechanism of action of SMARCAD1 and BRCA1 in maintaining replication fork integrity. SMARCAD1 maintains fork progression by regulating the PCNA occupancy at unperturbed replication forks to prevent fork stalling by blocking 53BP1 enrichment. While stalled forks require BRCA1-mediated fork protection when SMARCAD1 is off-loaded, efficient fork restart further requires SMARCAD1 to evict 53BP1 and restore PCNA levels by preventing PCNA unloading by ATAD5-RLC complex. The loss of 53BP1 can restore the PCNA levels, fork stability and genome stability in SMARCAD1-deficient cells. BRCA1 mediated fork protection against Mre11 DNA nuclease is essential to maintain fork progression in the SMARCAD1-deficient cells while SMARCAD1 is essential to maintain fork progression in BRCA1-deficient cells to maintain genome stability and subsequently cell survival.

**Table S1.**
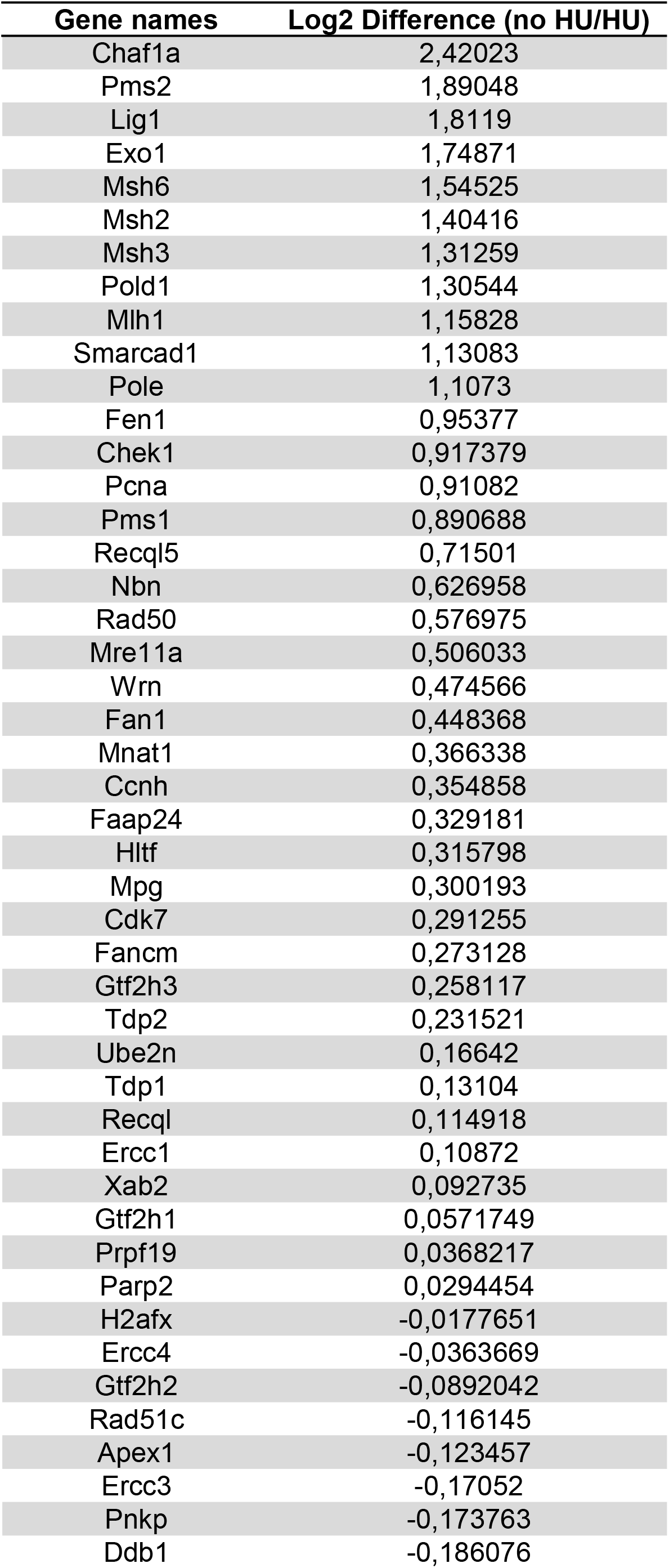
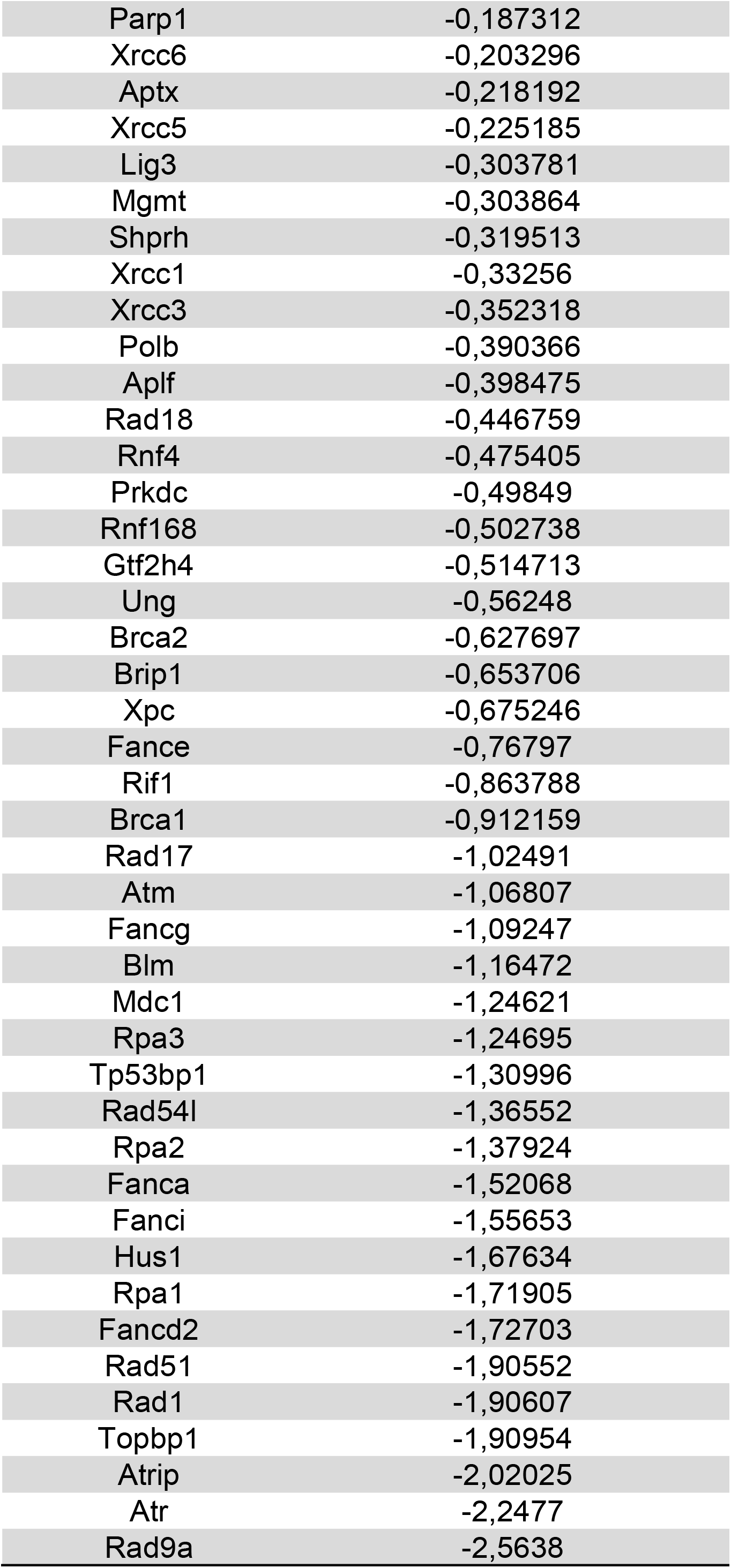
List of DDR proteins Enriched in unperturbed and HU treated conditions.

**Table S2:**
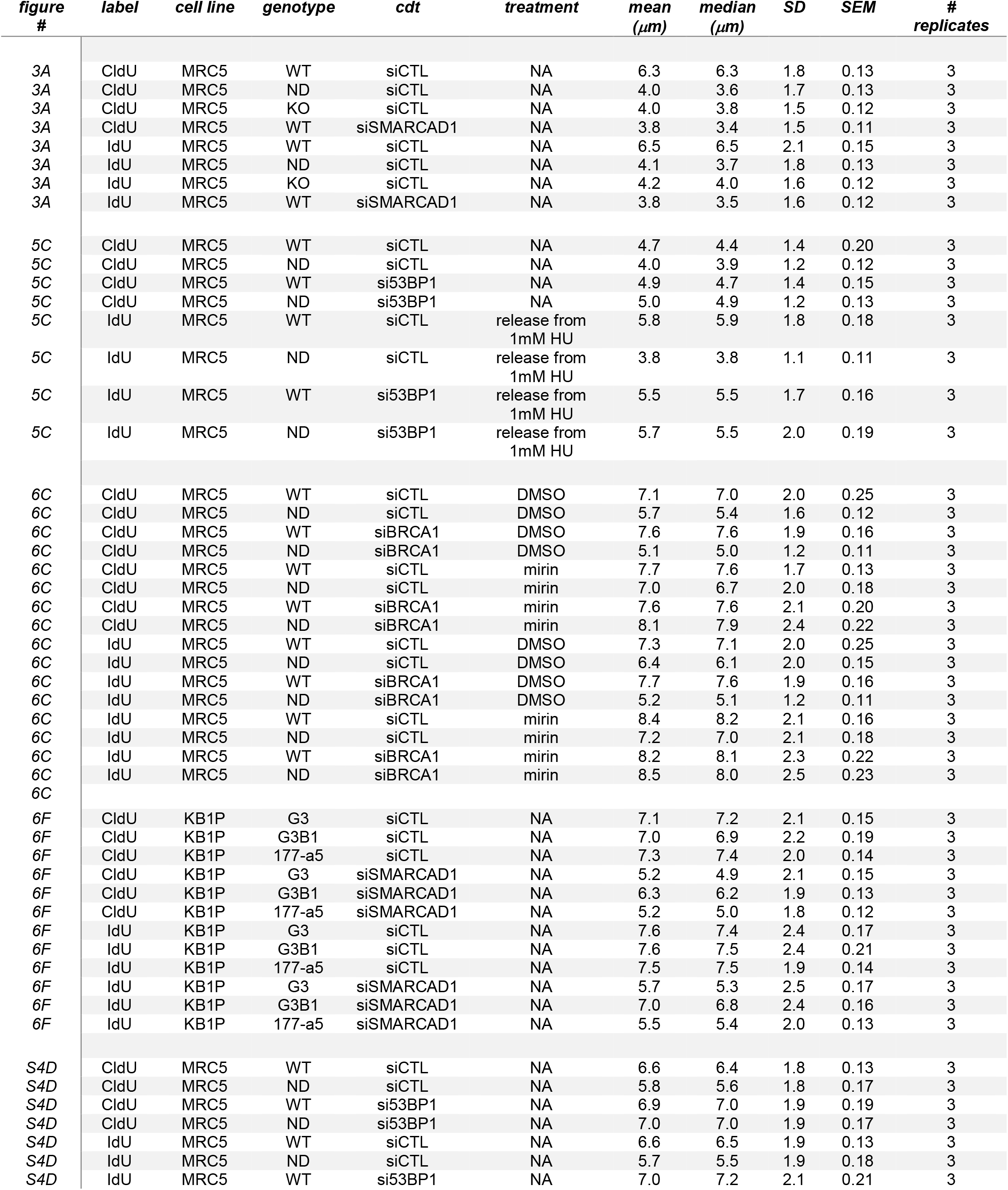
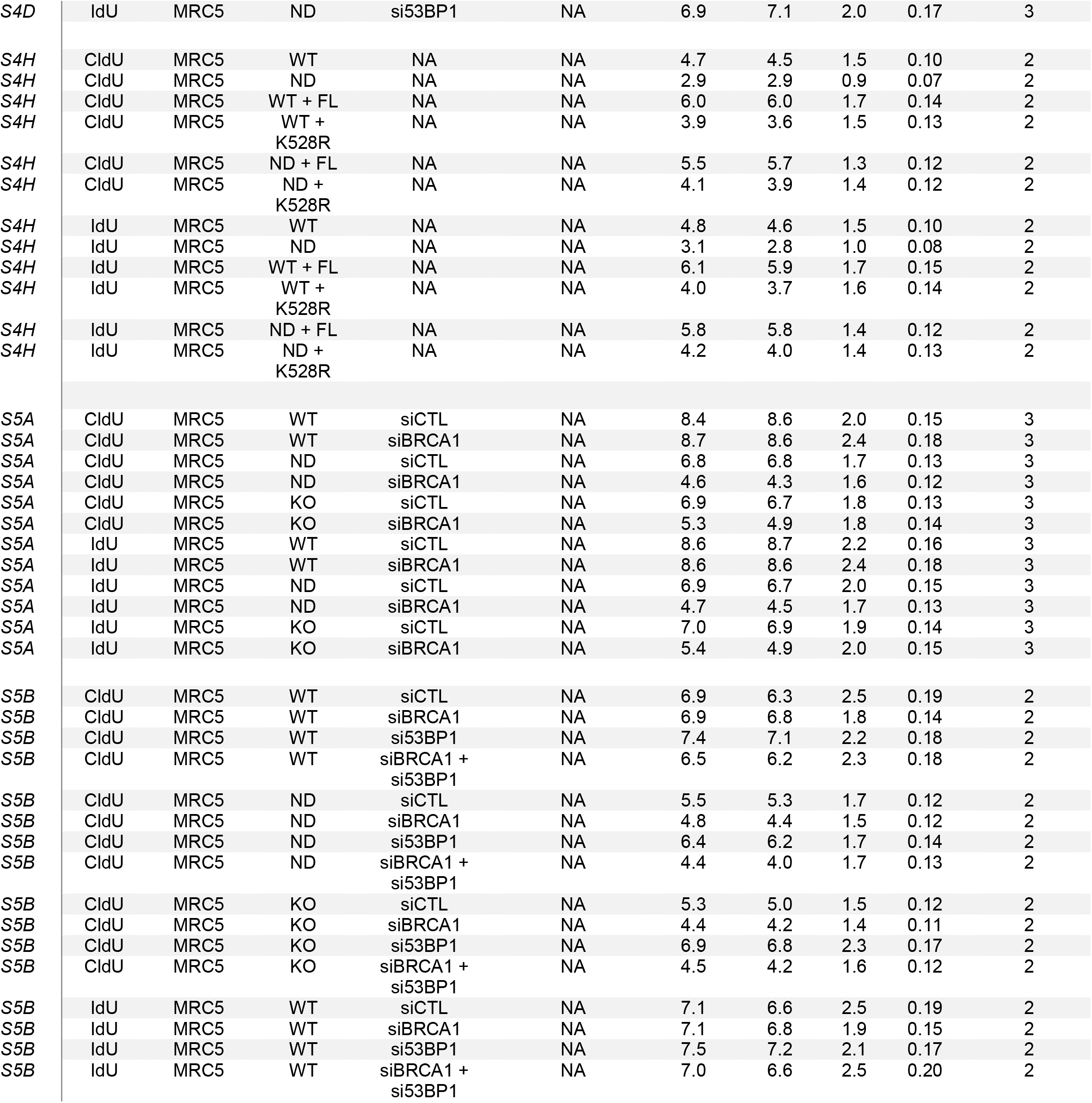
Summary of the DNA Fiber Spread Data Analysis. Mean, median, SD and SEM are the values for the plots shown in each respective figure. The number of experimental replicates is given in the column # replicates.

| figure # | label | cell line | genotype | cdt | treatment | mean (μm) | median (μm) | SD | SEM | # replicates |
| --- | --- | --- | --- | --- | --- | --- | --- | --- | --- | --- |
| 3A | CldU | MRC5 | WT | siCTL | NA | 6.3 | 6.3 | 1.8 | 0.13 | 3 |
| 3A | CldU | MRC5 | ND | siCTL | NA | 4.0 | 3.6 | 1.7 | 0.13 | 3 |
| 3A | CldU | MRC5 | KO | siCTL | NA | 4.0 | 3.8 | 1.5 | 0.12 | 3 |
| 3A | CldU | MRC5 | WT | siSMARCAD1 | NA | 3.8 | 3.4 | 1.5 | 0.11 | 3 |
| 3A | IdU | MRC5 | WT | siCTL | NA | 6.5 | 6.5 | 2.1 | 0.15 | 3 |
| 3A | IdU | MRC5 | ND | siCTL | NA | 4.1 | 3.7 | 1.8 | 0.13 | 3 |
| 3A | IdU | MRC5 | KO | siCTL | NA | 4.2 | 4.0 | 1.6 | 0.12 | 3 |
| 3A | IdU | MRC5 | WT | siSMARCAD1 | NA | 3.8 | 3.5 | 1.6 | 0.12 | 3 |
| 5C | CldU | MRC5 | WT | siCTL | NA | 4.7 | 4.4 | 1.4 | 0.20 | 3 |
| 5C | CldU | MRC5 | ND | siCTL | NA | 4.0 | 3.9 | 1.2 | 0.12 | 3 |
| 5C | CldU | MRC5 | WT | si53BP1 | NA | 4.9 | 4.7 | 1.4 | 0.15 | 3 |
| 5C | CldU | MRC5 | ND | si53BP1 | NA | 5.0 | 4.9 | 1.2 | 0.13 | 3 |
| 5C | IdU | MRC5 | WT | siCTL | release from 1mM HU | 5.8 | 5.9 | 1.8 | 0.18 | 3 |
| 5C | IdU | MRC5 | ND | siCTL | release from 1mM HU | 3.8 | 3.8 | 1.1 | 0.11 | 3 |
| 5C | IdU | MRC5 | WT | si53BP1 | release from 1mM HU | 5.5 | 5.5 | 1.7 | 0.16 | 3 |
| 5C | IdU | MRC5 | ND | si53BP1 | release from 1mM HU | 5.7 | 5.5 | 2.0 | 0.19 | 3 |
| 6C | CldU | MRC5 | WT | siCTL | DMSO | 7.1 | 7.0 | 2.0 | 0.25 | 3 |
| 6C | CldU | MRC5 | ND | siCTL | DMSO | 5.7 | 5.4 | 1.6 | 0.12 | 3 |
| 6C | CldU | MRC5 | WT | siBRCA1 | DMSO | 7.6 | 7.6 | 1.9 | 0.16 | 3 |
| 6C | CldU | MRC5 | ND | siBRCA1 | DMSO | 5.1 | 5.0 | 1.2 | 0.11 | 3 |
| 6C | CldU | MRC5 | WT | siCTL | mirin | 7.7 | 7.6 | 1.7 | 0.13 | 3 |
| 6C | CldU | MRC5 | ND | siCTL | mirin | 7.0 | 6.7 | 2.0 | 0.18 | 3 |
| 6C | CldU | MRC5 | WT | siBRCA1 | mirin | 7.6 | 7.6 | 2.1 | 0.20 | 3 |
| 6C | CldU | MRC5 | ND | siBRCA1 | mirin | 8.1 | 7.9 | 2.4 | 0.22 | 3 |
| 6C | IdU | MRC5 | WT | siCTL | DMSO | 7.3 | 7.1 | 2.0 | 0.25 | 3 |
| 6C | IdU | MRC5 | ND | siCTL | DMSO | 6.4 | 6.1 | 2.0 | 0.15 | 3 |
| 6C | IdU | MRC5 | WT | siBRCA1 | DMSO | 7.7 | 7.6 | 1.9 | 0.16 | 3 |
| 6C | IdU | MRC5 | ND | siBRCA1 | DMSO | 5.2 | 5.1 | 1.2 | 0.11 | 3 |
| 6C | IdU | MRC5 | WT | siCTL | mirin | 8.4 | 8.2 | 2.1 | 0.16 | 3 |
| 6C | IdU | MRC5 | ND | siCTL | mirin | 7.2 | 7.0 | 2.1 | 0.18 | 3 |
| 6C | IdU | MRC5 | WT | siBRCA1 | mirin | 8.2 | 8.1 | 2.3 | 0.22 | 3 |
| 6C | IdU | MRC5 | ND | siBRCA1 | mirin | 8.5 | 8.0 | 2.5 | 0.23 | 3 |
| 6C |  |  |  |  |  |  |  |  |  |  |
| 6F | CldU | KB1P | G3 | siCTL | NA | 7.1 | 7.2 | 2.1 | 0.15 | 3 |
| 6F | CldU | KB1P | G3B1 | siCTL | NA | 7.0 | 6.9 | 2.2 | 0.19 | 3 |
| 6F | CldU | KB1P | 177-a5 | siCTL | NA | 7.3 | 7.4 | 2.0 | 0.14 | 3 |
| 6F | CldU | KB1P | G3 | siSMARCAD1 | NA | 5.2 | 4.9 | 2.1 | 0.15 | 3 |
| 6F | CldU | KB1P | G3B1 | siSMARCAD1 | NA | 6.3 | 6.2 | 1.9 | 0.13 | 3 |
| 6F | CldU | KB1P | 177-a5 | siSMARCAD1 | NA | 5.2 | 5.0 | 1.8 | 0.12 | 3 |
| 6F | IdU | KB1P | G3 | siCTL | NA | 7.6 | 7.4 | 2.4 | 0.17 | 3 |
| 6F | IdU | KB1P | G3B1 | siCTL | NA | 7.6 | 7.5 | 2.4 | 0.21 | 3 |
| 6F | IdU | KB1P | 177-a5 | siCTL | NA | 7.5 | 7.5 | 1.9 | 0.14 | 3 |
| 6F | IdU | KB1P | G3 | siSMARCAD1 | NA | 5.7 | 5.3 | 2.5 | 0.17 | 3 |
| 6F | IdU | KB1P | G3B1 | siSMARCAD1 | NA | 7.0 | 6.8 | 2.4 | 0.16 | 3 |
| 6F | IdU | KB1P | 177-a5 | siSMARCAD1 | NA | 5.5 | 5.4 | 2.0 | 0.13 | 3 |
| S4D | CldU | MRC5 | WT | siCTL | NA | 6.6 | 6.4 | 1.8 | 0.13 | 3 |
| S4D | CldU | MRC5 | ND | siCTL | NA | 5.8 | 5.6 | 1.8 | 0.17 | 3 |
| S4D | CldU | MRC5 | WT | si53BP1 | NA | 6.9 | 7.0 | 1.9 | 0.19 | 3 |
| S4D | CldU | MRC5 | ND | si53BP1 | NA | 7.0 | 7.0 | 1.9 | 0.17 | 3 |
| S4D | IdU | MRC5 | WT | siCTL | NA | 6.6 | 6.5 | 1.9 | 0.13 | 3 |
| S4D | IdU | MRC5 | ND | siCTL | NA | 5.7 | 5.5 | 1.9 | 0.18 | 3 |
| S4D | IdU | MRC5 | WT | si53BP1 | NA | 7.0 | 7.2 | 2.1 | 0.21 | 3 |
| S4D | IdU | MRC5 | ND | si53BP1 | NA | 6.9 | 7.1 | 2.0 | 0.17 | 3 |
| S4H | CldU | MRC5 | WT | NA | NA | 4.7 | 4.5 | 1.5 | 0.10 | 2 |
| S4H | CldU | MRC5 | ND | NA | NA | 2.9 | 2.9 | 0.9 | 0.07 | 2 |
| S4H | CldU | MRC5 | WT + FL | NA | NA | 6.0 | 6.0 | 1.7 | 0.14 | 2 |
| S4H | CldU | MRC5 | WT + K528R | NA | NA | 3.9 | 3.6 | 1.5 | 0.13 | 2 |
| S4H | CldU | MRC5 | ND + FL | NA | NA | 5.5 | 5.7 | 1.3 | 0.12 | 2 |
| S4H | CldU | MRC5 | ND + K528R | NA | NA | 4.1 | 3.9 | 1.4 | 0.12 | 2 |
| S4H | IdU | MRC5 | WT | NA | NA | 4.8 | 4.6 | 1.5 | 0.10 | 2 |
| S4H | IdU | MRC5 | ND | NA | NA | 3.1 | 2.8 | 1.0 | 0.08 | 2 |
| S4H | IdU | MRC5 | WT + FL | NA | NA | 6.1 | 5.9 | 1.7 | 0.15 | 2 |
| S4H | IdU | MRC5 | WT + K528R | NA | NA | 4.0 | 3.7 | 1.6 | 0.14 | 2 |
| S4H | IdU | MRC5 | ND + FL | NA | NA | 5.8 | 5.8 | 1.4 | 0.12 | 2 |
| S4H | IdU | MRC5 | ND + K528R | NA | NA | 4.2 | 4.0 | 1.4 | 0.13 | 2 |
| S5A | CldU | MRC5 | WT | siCTL | NA | 8.4 | 8.6 | 2.0 | 0.15 | 3 |
| S5A | CldU | MRC5 | WT | siBRCA1 | NA | 8.7 | 8.6 | 2.4 | 0.18 | 3 |
| S5A | CldU | MRC5 | ND | siCTL | NA | 6.8 | 6.8 | 1.7 | 0.13 | 3 |
| S5A | CldU | MRC5 | ND | siBRCA1 | NA | 4.6 | 4.3 | 1.6 | 0.12 | 3 |
| S5A | CldU | MRC5 | KO | siCTL | NA | 6.9 | 6.7 | 1.8 | 0.13 | 3 |
| S5A | CldU | MRC5 | KO | siBRCA1 | NA | 5.3 | 4.9 | 1.8 | 0.14 | 3 |
| S5A | IdU | MRC5 | WT | siCTL | NA | 8.6 | 8.7 | 2.2 | 0.16 | 3 |
| S5A | IdU | MRC5 | WT | siBRCA1 | NA | 8.6 | 8.6 | 2.4 | 0.18 | 3 |
| S5A | IdU | MRC5 | ND | siCTL | NA | 6.9 | 6.7 | 2.0 | 0.15 | 3 |
| S5A | IdU | MRC5 | ND | siBRCA1 | NA | 4.7 | 4.5 | 1.7 | 0.13 | 3 |
| S5A | IdU | MRC5 | KO | siCTL | NA | 7.0 | 6.9 | 1.9 | 0.14 | 3 |
| S5A | IdU | MRC5 | KO | siBRCA1 | NA | 5.4 | 4.9 | 2.0 | 0.15 | 3 |
| S5B | CldU | MRC5 | WT | siCTL | NA | 6.9 | 6.3 | 2.5 | 0.19 | 2 |
| S5B | CldU | MRC5 | WT | siBRCA1 | NA | 6.9 | 6.8 | 1.8 | 0.14 | 2 |
| S5B | CldU | MRC5 | WT | si53BP1 | NA | 7.4 | 7.1 | 2.2 | 0.18 | 2 |
| S5B | CldU | MRC5 | WT | siBRCA1 + si53BP1 | NA | 6.5 | 6.2 | 2.3 | 0.18 | 2 |
| S5B | CldU | MRC5 | ND | siCTL | NA | 5.5 | 5.3 | 1.7 | 0.12 | 2 |
| S5B | CldU | MRC5 | ND | siBRCA1 | NA | 4.8 | 4.4 | 1.5 | 0.12 | 2 |
| S5B | CldU | MRC5 | ND | si53BP1 | NA | 6.4 | 6.2 | 1.7 | 0.14 | 2 |
| S5B | CldU | MRC5 | ND | siBRCA1 + si53BP1 | NA | 4.4 | 4.0 | 1.7 | 0.13 | 2 |
| S5B | CldU | MRC5 | KO | siCTL | NA | 5.3 | 5.0 | 1.5 | 0.12 | 2 |
| S5B | CldU | MRC5 | KO | siBRCA1 | NA | 4.4 | 4.2 | 1.4 | 0.11 | 2 |
| S5B | CldU | MRC5 | KO | si53BP1 | NA | 6.9 | 6.8 | 2.3 | 0.17 | 2 |
| S5B | CldU | MRC5 | KO | siBRCA1 + si53BP1 | NA | 4.5 | 4.2 | 1.6 | 0.12 | 2 |
| S5B | IdU | MRC5 | WT | siCTL | NA | 7.1 | 6.6 | 2.5 | 0.19 | 2 |
| S5B | IdU | MRC5 | WT | siBRCA1 | NA | 7.1 | 6.8 | 1.9 | 0.15 | 2 |
| S5B | IdU | MRC5 | WT | si53BP1 | NA | 7.5 | 7.2 | 2.1 | 0.17 | 2 |
| S5B | IdU | MRC5 | WT | siBRCA1 + si53BP1 | NA | 7.0 | 6.6 | 2.5 | 0.20 | 2 |

**Table S3.**
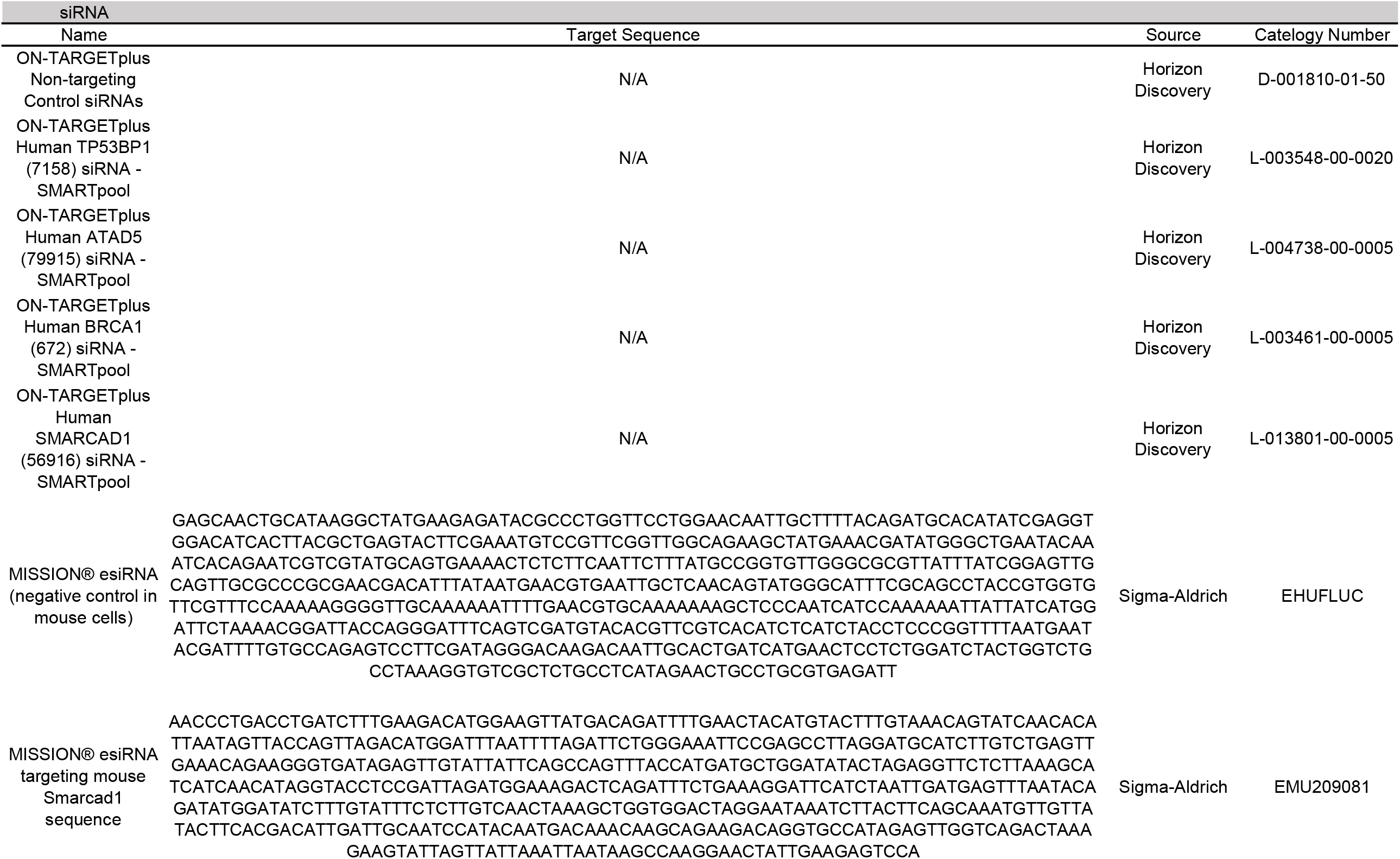
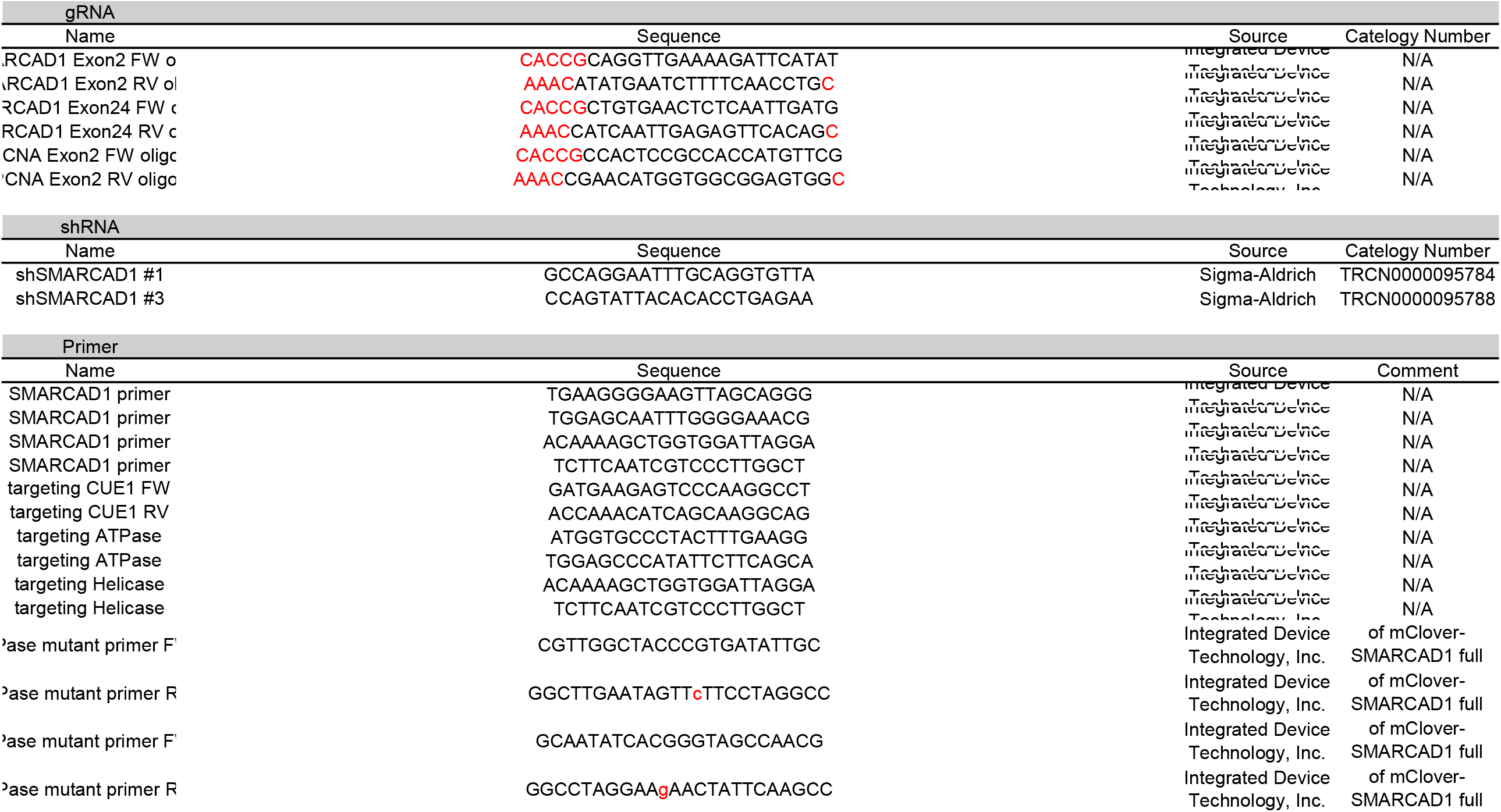
siRNA, gRNA, shRNA and primers used in this study.

## Notes

### Competing Interest Statement

The authors have declared no competing interest.

